# Functional states of prelimbic and related circuits during the acquisition of a GO/noGO task in rats

**DOI:** 10.1101/2024.03.25.586558

**Authors:** Carmen Muñoz-Redondo, Gloria G. Parras, Celia Andreu-Sánchez, Miguel Ángel Martín-Pascual, José M. Delgado-García, Agnès Gruart

**Affiliations:** Division of Neurosciences, Pablo de Olavide University, Seville, Spain; Neuro-Com Research Group, Department of Audiovisual Communication and Advertising, Universitat Autònoma de Barcelona, Barcelona, Spain; Research and Development, Institute of Spanish Public Television (RTVE), Corporación Radio Televisión Española, Barcelona, Spain

**Keywords:** basolateral amygdala, GO/noGO test, instrumental learning, nucleus accumbens septi, prelimbic cortex, primary motor cortex, rats, spectral analysis, touch screen

## Abstract

GO/noGO tasks enable assessing decision-making processes and the ability to suppress a specific action according to the context. Here, rats had to discriminate between two visual stimuli (GO or noGO) shown on an iPad screen. The execution (for GO) or non-execution (for noGO) of the selected action (to touch or not the visual display) were reinforced with food. The main goal was to record and to analyze local field potentials (LFPs) collected from cortical and subcortical structures when the visual stimuli were shown on the touch screen and during the subsequent activities. Rats were implanted with recording electrodes in the prelimbic cortex, primary motor cortex, nucleus accumbens septi, basolateral amygdala, dorsolateral and dorsomedial striatum, hippocampal CA1, and mediodorsal thalamic nucleus. Spectral analyses of the collected data demonstrate that the prelimbic cortex was selectively involved in the cognitive and motivational processing of the learning task but not in the execution of reward-directed behaviors. In addition, the other recorded structures presented specific tendencies to be involved in these two types of brain activity in response to the presentation of GO or noGO stimuli. Spectral analyses, spectrograms, and coherence between the recorded brain areas indicate their specific involvement in GO vs. noGO tasks.

## INTRODUCTION

Many experimental studies have convincingly shown that the medial prefrontal cortex (mPFC) represents one of the highest cortical centers dealing with goal-directed actions, decision-making tasks, associative learning, social interactions, and the execution of distinct types of adaptive behavior (Haroush and Williams, 2015; Lee et al., 2016; Carlén, 2017; Conde-Moro et al., 2019). While the electrical stimulation of the mPFC evokes behavioral freezing and prevents the acquisition of classical and instrumental conditioning tasks (Leal-Campanario et al., 2007; Jurado-Parras et al., 2012), the experimental lesion of the same areas modifies anxiety-behaviors and social interactions in rodents (Jinks and McGregor, 1997; Shah and Treit, 2003). The mPFC presents reciprocal connections with the core of the nucleus accumbens septi (NAc) and with the basolateral amygdala (BLA), two brain sites particularly related to positive and negative rewards, respectively (Vertes et al., 2004). Specifically, the prelimbic (PrL) and infralimbic cortices have differential projections to these two areas and present different functional capabilities in relation to cooperative learning, social interactions, active avoidance, and conditioned fear (Vidal-Gonzalez et al., 2006; Minami et al., 2017; Conde-Moro et al., 2019; Capuzzo and Floresco, 2020). In a similar way, the mPFC presents greater neural connections with the dorsomedial striatum (DMS)―a striatal area mostly related with the motivational control of goal-directed behaviors―than with the dorsolateral striatum (DLS), another striatal area mainly related with the elaboration of motor plans in accordance with its inputs from the motor and sensorimotor cortices (Stalnaker et al., 2010; Graybiel and Grafton, 2015; Corbit and Janak, 2016; Vandaele et al., 2019). The mPFC has reciprocal connectivity with the thalamic mediodorsal nucleus (MD) related to attentional states and working memory maintenance (Bolkan et al., 2017), and projects to the ipsilateral primary motor cortex (MC1) in relation to the final integration of ongoing motor commands and the acquisition and proper performance of new motor abilities (Kaufman et al., 2015; Wu et al., 2022). Finally, the mPFC presents interesting functional interactions with the hippocampal CA1 area (CA1) across the subiculum and the thalamic reuniens nucleus (Vertes et al., 2004; Eleore et al., 2011; Jurado-Parras et al., 2012; Harvey et al., 2023).

GO/noGO tests have many different uses in business, military, clinical, pharmacological, and psychological studies. More specifically, these tests have already been used for the study of different aspects of associative learning mechanisms (van Wingerden et al., 2014; Sicre et al., 2020), object recognition memories (Cole et al., 2020), and spatiotemporal activity patterns (Villa et al., 1999). We used this test to discriminate instrumentally conditioned responses involving or not body movements (such as approaching a lever to press it in order to collect a positive reward). The aim was to separate the acquisition of cognitive vs. purely behavioral functions. While the GO correct response involves approaching and touching a selected stimulus in order to collect a reward, response inhibitions (the noGO correct response) refer to the capability to suppress inappropriate responses in accordance with the learning situation (Bari and Robbins, 2013). It is generally accepted that these two executive functions are implemented by the prefrontal cortex (Chikazoe, 2010).

For the present study, we selected the PrL cortex because it is the mPFC area most directly related to instrumental conditioning, decision-making tasks, and related appetitive and consummatory behaviors (Hernández-González et al., 2017; Minami et al., 2017; Rocha-Almeida et al, 2018; Conde-Moro et al., 2019). We recorded LFPs generated in this cortical area and their relationships with the above-mentioned brain sites (NAc, BLA, DMS, DLS, MD, MC1, and CA1) during the performance of a GO/noGO test. In the case of the GO task, the experimental rat has to learn to associate a goal-directed action (to touch a horizontal white rectangular stimulus presented on an iPad screen) to collect a reinforcing stimulus (a pellet of food). In contrast, in the case of the noGO task, the animal has to inhibit the goal-directed behavior when a vertical green rectangular stimulus is presented on the iPad screen in order to collect the food reward. Thus, during the noGO task the animal has to actively suppress the goal-directed behavior. The multiple LFP recordings carried out in the above-mentioned cortical and subcortical areas provided important information with regard to their specific involvement, brain-state dynamics, and intrinsic coherences during the performance of a GO/noGO test, enabling a separation between cognitive vs. behavioral/motor components of the animal’s decisions to deal with significant external stimuli.

## METHODS

### Experimental animals

These experiments were carried out with male Lister Hooded rats provided by an authorized supplier (Charles Riber Laboratories, Barcelona, Spain). Rats were 3-4 months old and weighed 250-300 g at the beginning of the experiments. Following their arrival at the Pablo de Olavide Animal Facilities (Seville, Spain), animals were housed in individual cages until the end of the experiments. They were kept on a 12 h light/dark cycle in a stable environment of temperature (21 ± 1 °C) and humidity (55 ± 5%). Unless otherwise indicated (see below), animals had water and food available *ad libitum*.

### Ethical statement

All the experiments were performed in accordance with the regulations of the European Union Council (2010/276:33-79/EU), Spanish authorities (BOE 34:11370-11421, 2013), and ARRIVE guidelines (https://arriveguidelines.org) for the use of laboratory animals in chronic studies. Experiments were also approved by the corresponding Ethics Committees of Pablo de Olavide University and the Junta de Andalucía, Spain (codes 06/03/2018/025, 06/04/2020/049 and 10/10/2023/087).

### Surgical procedures

Following previously described surgical procedures (Hernández-González et al., 2017; Conde-Moro et al., 2019), animals were deeply anesthetized with 2% isoflurane delivered via an anesthesia mask (David Kopf Instruments, Tujunga, CA, USA). The anesthetic gas was supplied from a calibrated vaporizer (Fluotec-5, Fluotec-Olmeda, Tewksbury, MA, USA), at a flow rate of 1-2 L/min of oxygen (AstraZeneca, Madrid, Spain). Prior to electrode implantation, the rat was placed in a stereotaxic device (David Kopf Instruments). As illustrated in Fig. 1A-C, and in accordance with the Paxinos and Watson atlas (2007), animals were implanted with recording electrodes aimed at the following brain sites: i) anteroposterior from Bregma (AP): 3.72 mm; left (L): 3.4 mm; and depth from brain surface (De): 1.6 mm for the MC1. ii) AP: 3.24 mm; right (R): 0.5 mm; De: 2.5 mm for the PrL cortex. iii) AP: 2.04 mm; R: 1.5 mm; De: 6.3 mm for the NAc core. iv) AP: 0.6 mm; R: 3.8 mm; De: 4 mm for the DLS. v) AP: −0.36 mm; R: 2.6 mm; De: 4 mm for the DMS. vi) AP: −2.28 mm; R: 5 mm; De: 7.4 mm for the BLA. vii) AP: −3.2 mm; L: 0.8 mm; De: 5.2 mm for the MD. And viii) AP: −4.56 mm; L: 2.2 mm; De: 2.2 mm for the CA1. All of these electrodes were handmade from 50 µm tungsten wire coated with Teflon (Advent Research, Eynsham, UK). Each recording electrode consisted of two tungsten wires with a separation at the tips of ≍ 0.2 mm. For better surface exposure, Teflon coating was removed from electrode tips ≍ 0.1 mm. To serve as a ground, animals were also implanted with two bare silver wires (0.1 mm in diameter) affixed to the skull with small screws. Electrodes were soldered to two 6-pin sockets (RS-Amidata, Madrid, Spain). The two sockets were fixed to the skull with some additional small screws and dental cement. Animals were allowed a week to recover from surgery before the start of the experimental sessions.

**Figure 1.**
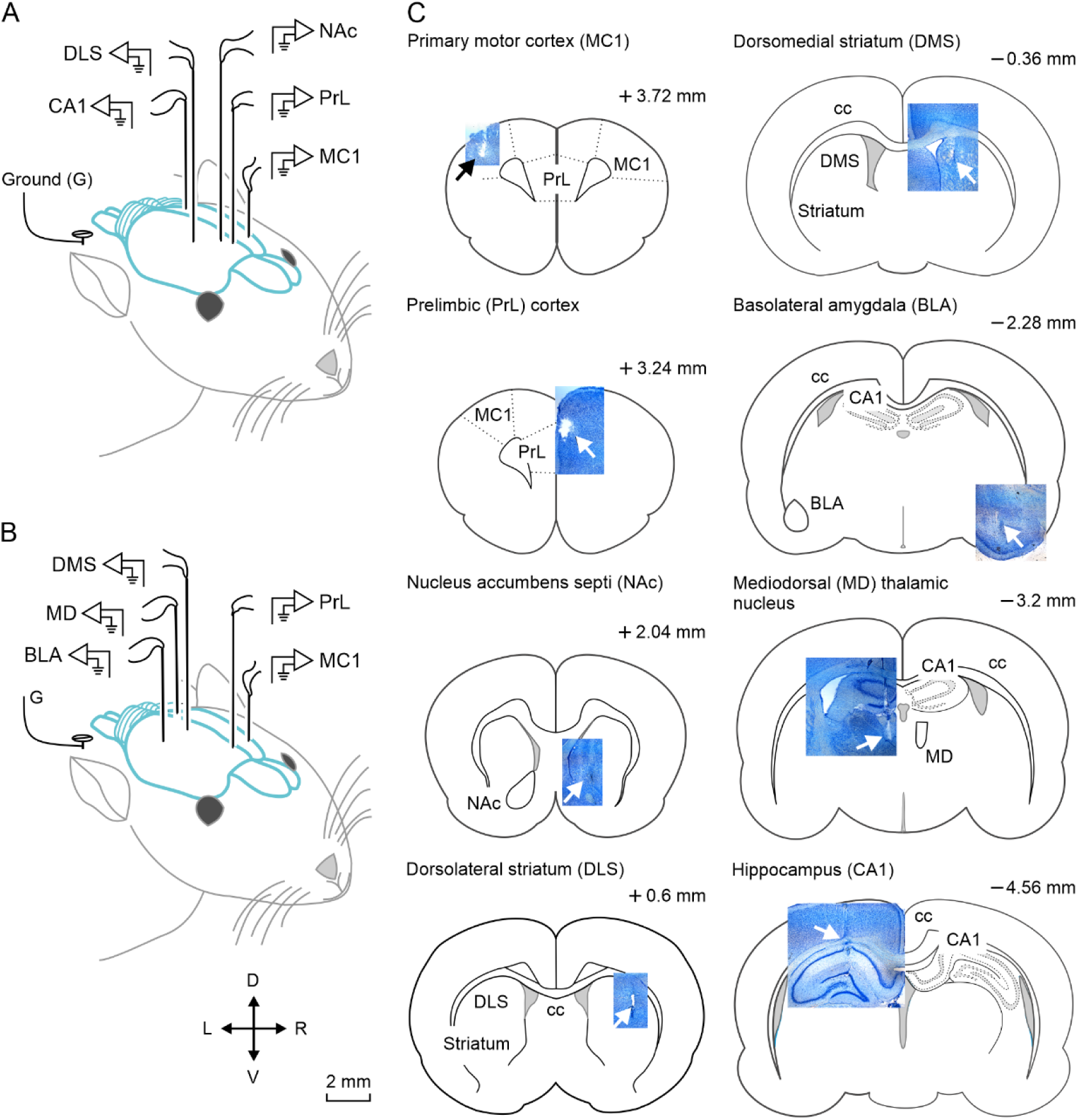
Experimental design for LFP recordings. **A, B.** Schematic representations of the location of recording electrodes. Half of the rats (n = 5) were implanted with pairs of recording electrodes in the right prelimbic (PrL) cortex, the core of the nucleus accumbens septi (NAc) and the dorsolateral striatum (DLS), as well as in the left primary motor cortex (MC1) and the dorsal hippocampal CA1 area (CA1) (**A**). The other half (n = 5) of the animals were implanted with pairs of recording electrodes in the right PrL cortex, the dorsomedial striatum (DMS) and the basolateral amygdala (BLA), as well as in the left MC1 and the mediodorsal thalamic nucleus (MD) (**B**). **C**. Diagrams of brain sagittal sections including representative photomicrographs illustrating the final location of implanted electrodes (arrows). The rostrocaudal coordinates of illustrated brain sections are indicated (Paxinos and Watson, 2007). Abbreviations: cc. corpus callosum; D, dorsal; L, left; R, right; V, ventral.

### Implementation of GO/noGO program on the touch screen

To implement the GO/noGO program, we selected the touch screen procedure (Horner et al., 2013; Hernández-González et al., 2017). We used a conventional Skinner box modified by the addition of an iPad touch screen as one of the walls of the chamber. We used a capacitive multi-touch screen device, iPad 2 model, with a 9.7–inch LED–backlit glossy widescreen multi-touch display using the IPS technology (Apple Inc., Cupertino, CA, USA). The screen had 1024 × 768 pixel resolution at 132 pixels per inch. We developed an ad hoc software (RatButton 2.2. for iOS 8.4) for stimuli presentation on the touch screen. The RatButton software was developed with the X-Code in object-oriented programming language Objective-C. Stimulus color (sRGB IEC61966- 2.1: 42, 246, 42 for green and 255, 255, 255 for white) and luminance (150 cd/m^2^ at 20 cm distance) were adapted to rats’ vision (Jacobs et al., 2001) with the help of a colorimeter (Minolta Chroma Meter xy-1, Konica Minolta Inc., Marunouchi, Tokyo, Japan) and a photometer (Sekonic DualMaster L-558/L; Sekonic Corporation, Marunouchi, Tokyo, Japan).

To start the session, the screen was illuminated in red (Fig. 2B). Each trial (n = 40 per session) was signaled with a tone (5,700 Hz, 85 dB, 100 ms) 100 ms before the stimulus. For the GO stimulus we used a horizontal white rectangle with a black background, whereas as a noGO stimulus we used a vertical green rectangle with a black background. Stimuli were centered in the upper half of the screen, at 159-x and 100-y coordinates. Each colored rectangle (white for GO and green for noGO) measured 450 × 150 px (horizontal, GO) and 150 × 450 px (vertical, noGO). In both cases, the rectangles were inserted in identical-sized virtual touchable buttons of 450 × 450 px. During GO trials (Fig. 2C), the rat was reinforced with a pellet of food if it touched the white horizontal rectangle during the illuminated period (10 s to 5 s). In the case of noGO trials (Fig. 2D), the animal had to avoid touching the vertical green rectangle during the illuminated period (2.5 s to 5 s) in order to obtain a food pellet as a reward. The incorrect answer did not give any reward or punishment. Each trial had an inter-trial interval of 30 s.

**Figure 2.**
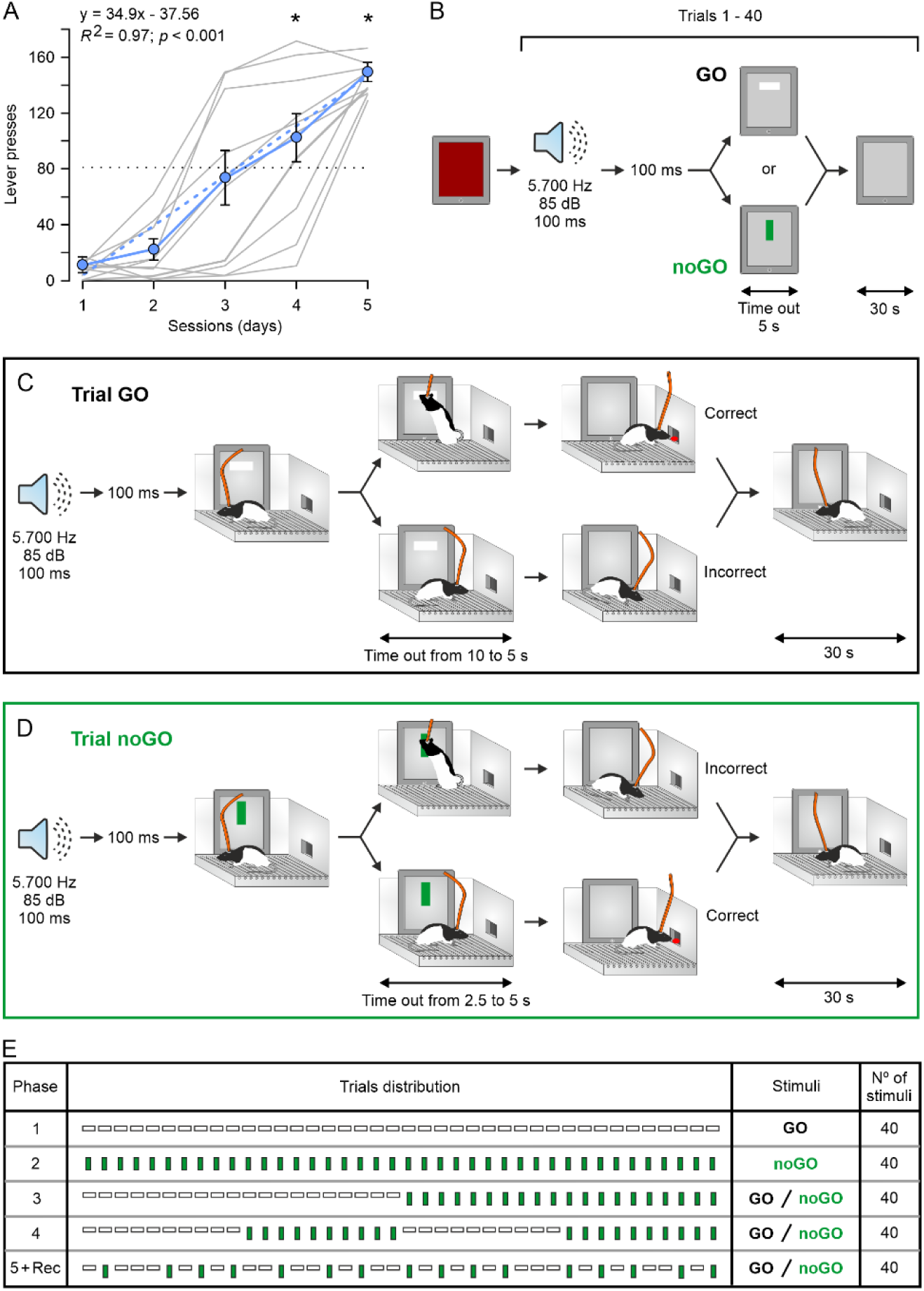
Experimental design for GO/noGO training. **A.** In a first experimental step, animals were trained to press a lever in order to collect a food pellet with a fixed-ratio (1:1) schedule. All animals (n = 10) reached criterion (to press the lever ≥ 80 times for two successive 20-minute session) by the 4th-5th conditioning session. Individual learning curves are illustrated, and the best linear adjustment is indicated with a dashed line. *, *p* < 0.05. **B-D**. General design for the GO/noGO task. Each session started with the iPad tablet illuminated in red. Stimuli for GO (a horizontal white rectangle on a black background) or noGO (a vertical green rectangle on the same black background) responses were presented after a preceding tone lasting for 100 ms. Independently of the correct or incorrect response, each trial lasted for 30 s. In accordance, each 20-minute session allowed the presentation of 40 trials. **E**. The whole experiment (5 phases) lasted 25 days. The duration of each phase depended upon the time to reach the selected criterion (see Fig. 3). Stimulus presentations during each phase are indicated. Local field potentials (LFPs) were recorded (Rec) and collected during the 5th phase exclusively. The duration of the GO stimulus was a maximum of 10 s for phase 1 and 5 s for phases 3 and 5, whereas the duration of the noGO stimulus was 2.5 s for phase 2 and 5 s for the following three phases.

### Instrumental conditioning in a conventional Skinner box

In a preliminary series of experiments (Fig. 2A), and in accordance with previous descriptions (Hernández-González et al., 2017), rats were trained for instrumental conditioning in a conventional (29.2 × 24.1 × 21 cm) Skinner box module (Med Associates, St. Albans, WT, USA). The operant conditioning box was housed within a soundproof container (90 × 55 × 60 cm) provided with a 45 dB white noise and a constantly illuminated (19 W) lamp (Cibertec, Madrid, Spain). The Skinner box was equipped with a lever and a food dispenser. The conditioning paradigm was programmed to deliver a pellet of food (Noyes formula P of 45 mg. Sandown Scientific, Hampton, England) every time the animal pressed the lever.

Before training in the Skinner box, rats were handled daily, and food-deprived until reaching ≍ 90% of their *ad libitum* feeding weight. Once the desired weight was achieved, animals were placed daily in the Skinner box for 20 min and allowed to press the lever to receiver a food pellet from the dispenser, using a fixed-ratio (1:1) schedule, until the selected criterion (to press the lever ≥ 80 times in each of two consecutive sessions) was reached (Hasan et al., 2013; Hernández-González et al., 2017).

Once accomplished the selected criterion (Fig. 2A), animals were transferred to the GO/noGO conditioning program. In this case, animals were placed in the above-described modified Skinner box in which one of the walls was substituted by a touch screen. Rats were allowed two habituation sessions before the GO/noGO test was started.

For phase 1 (Fig. 2E), only GO stimuli were available to receive a pellet of food delivered at the dispenser. The rat was supposed to touch the screen exactly in the area in which the stimulus (horizontal white rectangle) was presented. Each stimulus presentation was preceded by a 5,700 Hz tone lasting for 100 ms (Fig. 2C). The horizontal white rectangle remained illuminated for 10 s. When touched, the illumination disappeared, and a food pellet was delivered. Whether the screen was touched or not, the whole trial had a duration of 30 s, after which a new trial was started. A total of 40 trials presentations were carried out during the 20 min session. Criterion for this phase was to touch the screen for ≥ 60 times on each of two successive days (Fig. 3A,B).

**Figure 3.**
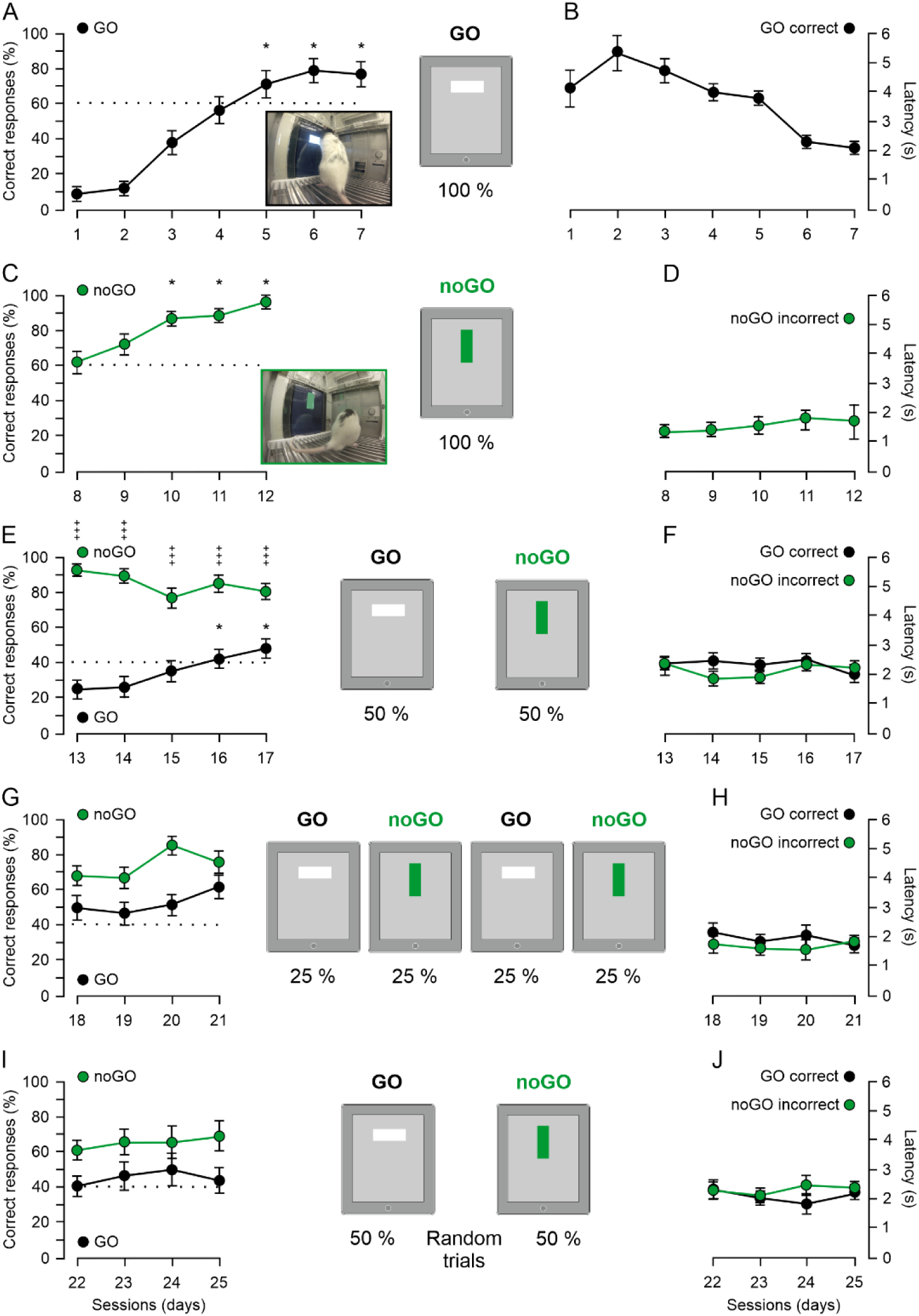
Validation of the GO/noGO test in rats that followed the 5 successive experimental phases. As illustrated in **A**, **C**, **E**, **G,** and **I,** the duration of each phase depended upon the time to reach the selected criterion (indicated by the dotted lines). Rats had to pass criterion for two successive sessions before moving to the next phase. Stimuli presented for each phase are illustrated. The two photographs illustrate stimuli for GO (**A**) and noGO (**B**) tasks, and a correct response of the representative animal. Note that the latency (**B**, **D**, **F**, **H** and **J**) for a correct response decreased across sessions to reach a minimum of ≍ 2.5 s during the final sessions. Although animals were connected to the recording system during all the recording sessions, LFPs were collected and analyzed only during the 5^th^ + Rec phase (**I**). The reason was that the unpredictable presentation of GO and noGO stimuli prevented a fast adaptation of the experimental animals to the stimulus sequence. *, *p* < 0.05; ^+++^, *p* < 0.001.

For phase 2 (Fig. 2E), only noGO stimuli were presented (Fig. 2D). As during phase 1, each stimulus presentation was preceded by a 100 ms tone. In this case, the noGO visual stimulus was presented for only 2.5 s. The duration of a complete trial was also 30 s, and the total number of trials was 40 for the 20 min session. Criterion was not to touch the vertical green rectangle for ≥ 60 times on each of two successive days (Fig. 3C,D).

Phase 3 (Fig. 2E) consisted of 20 trials presentations of the GO stimulus (the first half of the session) followed by another 20 trials presenting the noGO stimulus (the second half of the session). All the other parameters remained as described above, but the duration of both visual stimuli was 5 s. Criterion was to touch the screen for GO and not to touch it for noGO presentations ≥ 40 times on each of two successive days (Fig. 3E,F).

For phase 4, stimuli were presented in blocks of 10 trials each, following the sequence: 25% GO, 25% noGO, 25% GO, and 25% noGO (Fig. 2E). The duration of each stimulus was 5 s. All the other stimulus conditions remained as for phase 1. Criterion was as for phase 3: to touch the screen (for GO) and not to touch it (for noGO) ≥ 40 times on each of two successive days (Fig. 3G,H).

Finally, for phase 5 + Rec (recording), 20 stimuli for GO and another 20 for noGO were presented at random (Fig. 2E). Both GO and noGO stimuli lasted for 5 s. Criterion was as for phases 3 and 4. LFPs were collected during this phase (Fig. 3I,J).

LFPs were recorded using Grass P511 differential amplifiers with a bandwidth between 0.1 Hz and 3 kHz (Grass Telefactor, West Warwick, RI, USA). For the sake of homogeneity, and considering the difficulties on the selected training program, LFPs were recorded exclusively during the final experimental phase (5 + Rec; Fig. 2E).

### Histology

At the end of the experiments, and following standard procedures of our lab (Hernández-González et al., 2017; Conde-Moro et al., 2019), rats were deeply anesthetized with a saline solution of ketamine (100 mg/kg) and medetomidine (0.1 mg/kg) and perfused transcardially with 0.4% paraformaldehyde dissolved in phosphate-buffered saline (PBS, 0.1 M and 7.4 pH). Brains were cryoprotected with a PBS solution of 30% sucrose for a week before cutting them in coronal sections (50 µm) with the help of a sliding freezing microtome (Leica SM2000R, Nussloch, Germany). Sections including the implanted sites were mounted on gelatinized glass slides and stained, using the Nissl technique, with a PBS solution of 0.1% toluidine blue. After rinsing in PBS, sections were coverslipped and photographed using a Leica DM5000 B microscope (Leica Microsystems GmbH, Wetzlar, Germany) to determine the proper location of implanted electrodes (Fig. 1C).

### Data collection and analysis

Data for analysis and representations were collected from 10 animals (5 from group A and 5 from group B, see Fig. 1A,B) that completed the five experimental phases (Fig. 2E), reached all the expected criteria, and presented LFPs free of unwanted artifacts.

Conditioning programs, lever presses, screen touches, response latencies, delivered reinforcements, and recorded LFPs were monitored and acquired with the help of a MED-PC program (MED Associates). In addition, all the training sessions were recorded with a video capture system (Sony HDR-SR12E, Tokyo, Japan) synchronized to the operant conditioning program and to the LFP recordings. All these data were stored digitally on a computer through a CED 1401 Plus analog-to-digital converter (Cambridge Electronics Design, Cambridge, England). LFPs were sampled at 5 kHz with an amplitude resolution of 16 bits.

For frequency- (Figs. 5-7, 11) and time-domain (Figs. 8-10) analyses, we selected LFP epochs from the 2 s preceding screen touches during GO correct and noGO incorrect responses and the 2 s preceding the visual stimulus being switched off during noGO correct and GO incorrect responses. For the 8 recorded brain areas, a total of 20 samples (2 s each) collected from ≥ 5 rats were selected, analyzed, and represented in the above-mentioned figures. Frequency-domain analyses illustrated in Fig. 11 were carried out with 20-second samples of LFPs recorded before (sessions 1 and 2) and after (sessions 14 and 18) learning in the GO/noGO test for four selected (PrL, NAc, DLS, and CA1) brain areas. Analyses in the frequency domain were carried out according to the following bands: delta (1-6 Hz), theta (6-12 Hz), beta (12-30 Hz), low-gamma (30-50 Hz), and high-gamma (50-120 Hz). A high-pass filter (0-1 Hz) was applied to remove movement artifacts.

The Spike2 software (Cambridge Electronic Design) was used for the proper quantification of animal performance in the conventional and modified Skinner boxes. Following previous descriptions by some of us (Hernández-González et al., 2017; Conde-Moro et al., 2019), analytical tools for signal processing in the frequency domain (power spectra; Figs. 5-7 and 11), the multitaper Fourier transforms in the time-frequency domain (spectrograms, Figs. 8-9), and coherograms (Fig. 10) were developed from programs written in MATLAB (The MathWorks, Natick, MA, USA) and scripts of Chronux toolbook and software (Mitra and Bokil, 2008; version 2.12 Website: http://chronux.org/). Power spectra and spectrograms were computed using 5 tapers (K), and time-bandwidth (TW) of 3 [params.tapers = (TW = 3K = 5)]. A moving window of 0.5 s, shifted in 10-ms increments (movingwin = [0.5 0.01]), was also used for spectrogram analyses (Conde-Moro et al., 2019).

Collected results were processed for statistical analysis using the Sigmaplot 11.0 software (Systat Software Inc., San Jose, CA, USA). For Figs. 2 and 3A-D, we used the one-way repeated measures ANOVA or the Friedman repeated measures ANOVA on ranks. The Holm-Sidak method or the Tukey test were used to compare the different sessions when the main test was significant. For Fig. 3E-J, we used the two-way repeated measures ANOVA (with two-factor repetition). In cases that the F value was significant, we applied the Holm-Sidak method to find the differences between sessions and/or between stimuli (GO/noGO).

For Figs. 5-7, we applied the two-way repeated measures ANOVA followed by the Holm-Sidak method when needed. In Figs. 5-6, the spectral power of the four types of response was analyzed. In addition, in Fig. 7 the power spectral density (PSD) and the dominant frequency for the eight recording sites were studied.

Finally, for Fig. 11, we used the two-way repeated measures ANOVA to analyze LFP oscillations collected from four selected sites (PrL, NAc, DLS, and CA1). When needed, the Holm-Sidak method was used to find the statistical differences between the two factors―before and after learning, and at initial and final parts of the recording sessions.

In all of the cases, the significance level (*p* value) is indicated: *, *p* < 0.05; **, *p* < 0.01; and ***, *p* < 0.001. Data are always represented as the mean ± SEM.

## RESULTS

### Rats can learn a GO/noGO task, including the generation of active behavioral responses and the active inhibition of any motor response

As already mentioned in the Methods section and further illustrated in Fig. 1, rats were implanted with pairs of recording electrodes in the PrL cortex, MC1, NAc, DLS, and hippocampal CA1 area (Fig. 1A) or in the PrL, MC1, BLA, DMS, and MD (Fig. 1B). This electrode distribution avoided any putative damage to neural tissue caused by an excessive concentration of implanted electrodes. PrL and MC1 electrodes were implanted in both sets of rats for comparison with the other implanted electrodes. After the end of the experiments, we checked the correct location of implanted electrodes (Fig. 1C).

The final aim of the study was to determine the activity of LFPs recorded from all the implanted sites during the random presentation of GO or noGO stimuli in an instrumental conditioning task. A total of 5 rats for each electrode distribution (Fig. 1A,B) that correctly performed all the conditioning tasks and presented LFPs free of movement artifacts were selected. The following descriptions and analysis are centered on data collected from these 10 animals.

In a first experimental step, animals were placed in a conventional Skinner box in order to acquire an operant task consisting of pressing a lever in order to collect a piece of food, using a fixed-ratio (1:1) conditioning paradigm. Conditioning sessions lasted for 20 min and criterion was to press the lever for ≥ 80 times/session for two successive sessions. As illustrated in Fig. 2A, the 10 selected rats reached criterion by the 5th conditioning session (*F*_(4,36)_ = 37.091; *p* < 0.001). In this case, criterion was reached in the same sessions as (Fernández-Lamo et al., 2016) or earlier than (Hernández-González et al., 2017) previous reports using the same experimental set-up.

Once animals had correctly acquired this initial conditioning task, they were transferred to the GO/noGO conditioning program. Animals were placed in a modified Skinner box in which one of the walls was substituted by an iPad touch screen (Fig. 2B-D). Rats were allowed two habituation sessions before the GO/noGO task was started. When the rat was carefully placed in this modified Skinner box, the iPad screen was initially illuminated in red. For the GO response, the stimulus consisted of an illuminated horizontal white rectangle surrounded by a black background. The stimulus lasted for 10 to 5 s depending on the experimental phase (see below). The stimulus was removed as soon as it was touched by the rat or at the end of the programmed time (Fig. 2C). The stimulus for the noGO response consisted of a vertical green rectangle lasting from 2.5 to 5 s depending on the experimental phase. Here again, the stimulus was removed if touched or at the end of the programmed time (Fig. 2D). In both situations, the duration of each trial was 30 s. In this way, a maximum of 40 trials (stimuli) could be presented per session.

Instrumental conditioning of this GO/noGO task was divided into five successive phases (Fig. 2E). During phase 1, only stimuli corresponding to the GO task were presented (a total of 40 trials/session). Rats reached criterion (to touch the illuminated white rectangle in ≥ 60% of the trials) by the 7th conditioning session [*F*_(6,54)_ = 59.946; *p* < 0.001] (Fig. 3A). During this experimental phase, the visual stimulus lasted for 10 s, and the latency to touch the illuminated white rectangle decreased significantly [*F*_(6,54)_ = 14.727; *p* < 0.001] from ≍ 4 s during the 1st session to ≍ 2 s during the 7th session (Fig. 3B).

Through phase 2 (Fig. 2E), only stimuli corresponding to the noGO task were presented (also, a total of 40 trials/session). This task appeared to be easier to acquire for the experimental rats, as they reached criterion (to avoid touching the illuminated green rectangle in ≥ 60% of the trials) by the 4th conditioning session [χ^2^ = 28.557 with 4 degrees of freedom; *p* < 0.001] (Fig. 3C). During this phase, the visual stimulus was present for 5 s, and the latency for noGO incorrect responses was maintained for ≍ 1.5 s (Fig. 3D).

For phase 3 (Fig. 2E), sessions consisted of 20 presentations of the GO task followed by the same number of presentations (20) of the noGO task (a total of 40 presentations). Criterion was to generate ≍ 40% of GO and noGO correct responses. In this case, visual stimuli for GO and noGO tasks were present for a maximum of 5 s. This phase was significantly more difficult for the acquisition of the GO task vs. the noGO one [*F*_(4,36)_ = 6.800, *p* < 0.001] (Fig. 3E). Animals reached criterion for the GO task (to touch the illuminated white rectangle in ≥ 40% of the trials) by the 5th conditioning session (*t* = 4.679-3.186; *p* < 0.05). Latencies for touching the visual stimuli were maintained ≍ 2 s for correct GO responses and also for incorrect noGO responses (Fig. 3F).

During phase 4 (Fig. 2E), sessions consisted of 10 successive presentations of GO, noGO, GO, and noGO tasks (25% each). In this case, animals carried out a better performance of the GO tasks, and no significant differences were observed between GO and noGO correct responses [*F*_(3,27)_ = 2.849; *p* = 0.056] (Fig. 3G). The percentage of correct responses was over criterion for the four sessions and GO or noGO tasks. Latencies for correct GO and incorrect noGO responses were ≍ 2 s (Fig. 3H).

Finally, for phase 5 + Rec (Fig. 2E), sessions consisted of a random presentation of 20 GO and 20 noGO trials. At this stage, animals were perfectly trained to carry out the required task, reaching the selected criterion for the four conditioning sessions. No significant differences were observed in the percentage of correct GO vs. noGO responses [*F*_(3,27)_ = 0.410; *p* = 0.747]. Results presented in Fig. 3 were similar to those previously reported for male rats using a similar GO/noGO task (Zhang et al., 2023).

### Power spectra of five selected frequency bands from LFPs recorded in the 8 implanted areas were modified by the different experimental situations

As already indicated, LFPs were recorded from the 8 implanted areas during phase 5 + Rec from a total of 10 animals for four experimental sessions (see above). Fig. 4 is an illustration of LFPs recorded from 2 representative animals during the proper response to GO and noGO stimulus presentations. In the four cases, the dashed lines indicate the LFPs recorded during the presentation of the stimulus selected for GO (left) or noGO (right) expected responses. From the illustrated traces, the presence of delta/theta waves preceding the moment when the rat touched the horizontal white rectangle (left set of records) could be easily observed in different recorded areas (PrL, BLA, MD, DMS, and MC1) (Fig. 4A). For correct noGO responses, similar delta/theta waves could also be observed in different recorded areas (PrL, BLA, MD, DMS, NAc, and DLS) during the presentation of the vertical green rectangle (Fig. 4B).

**Figure 4.**
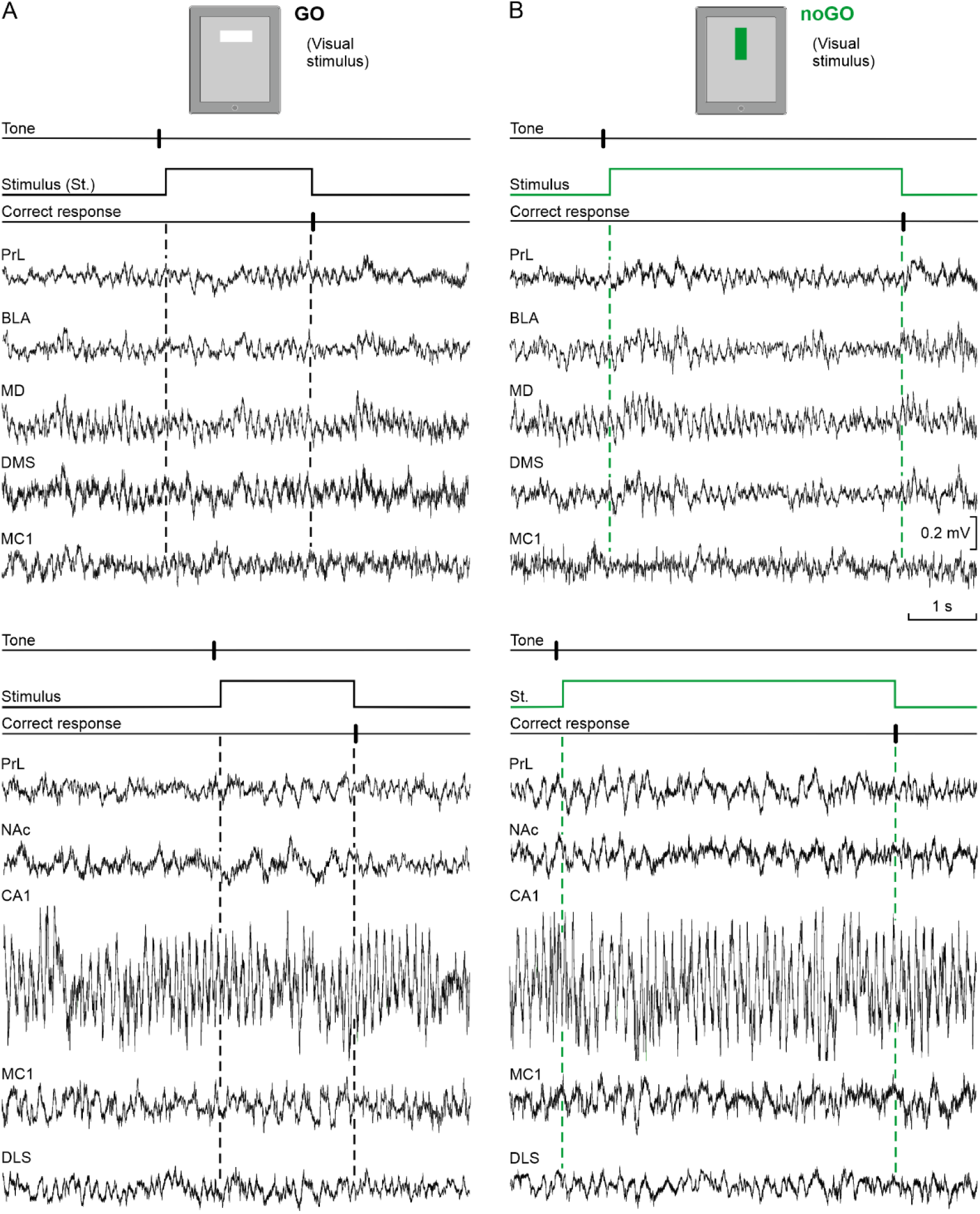
LFPs recorded from two representative animals implanted with recording electrodes as indicated in Fig. 1B (the top two sets of records) and Fig. 1A (the bottom two sets of records) during the performance of correct GO or noGO responses. **A, B**. From top to bottom are indicated the tone presentation, the presence of the stimulus for correct GO (**A**) and noGO (**B**) responses, the moment when the correct response was performed, and representative LFP oscillations collected from animals belonging to the B (top sets of records) or A (bottom sets of records) groups. LFP records were collected during the 5 + Rec phase. Note that for the GO condition (the left two sets of records) the two animals touched the iPad screen (horizontal white rectangle) in ≍ 2 s, while for the noGO condition (the right two sets of records) they waited the corresponding 5 s for the extinction of the stimulus (vertical green rectangle). Also note the presence of evident delta/theta oscillations just before the animal touched the screen in most of the illustrated traces (**A,** top and bottom). Interestingly, these delta/theta oscillations were also noticeable at the stimulus presentations during the noGO task (**B,** top and bottom).

As observed during the ongoing experiments and on the videotapes, animals were attentive to the visual stimuli presented on the iPad screen. In the case of GO correct and noGO incorrect responses, rats usually touched the screen in ≍ 2 s―i.e., well before the programed duration of the stimulus (5 s). In contrast, in the case of the noGO correct and GO incorrect responses, animals remained static until the end of the stimulus (5 s) when they moved directly to the feeder. Given those temporal differences, and for a detailed consideration of oscillatory activities presented during the four experimental situations, we selected 2-second LFP segments (a total of 20 from ≥ 5 animals) preceding GO correct responses and noGO incorrect responses, and from the 2 s preceding the moment when visual stimuli were switched off for noGO correct and GO incorrect responses.

In Figs. 5 and 6 are illustrated the spectral powers of 2-second segments selected from LFPs recorded during the four experimental situations. All these analyses were carried out using the fast Fourier transform method (see Methods). The power spectra (1-120 Hz) collected from the 8 selected areas are illustrated. Interestingly, both the CA1 and the MD presented greatest spectral powers for the theta band, while all the other recorded areas presented dominant peaks for the delta band. A quantification of mean spectral powers corresponding to the five selected bands (delta, theta, beta, and low- and high-gamma) included in the present study was also carried out (see insets of the five sets of histograms in Figs. 5 and 6).

**Figure 5.**
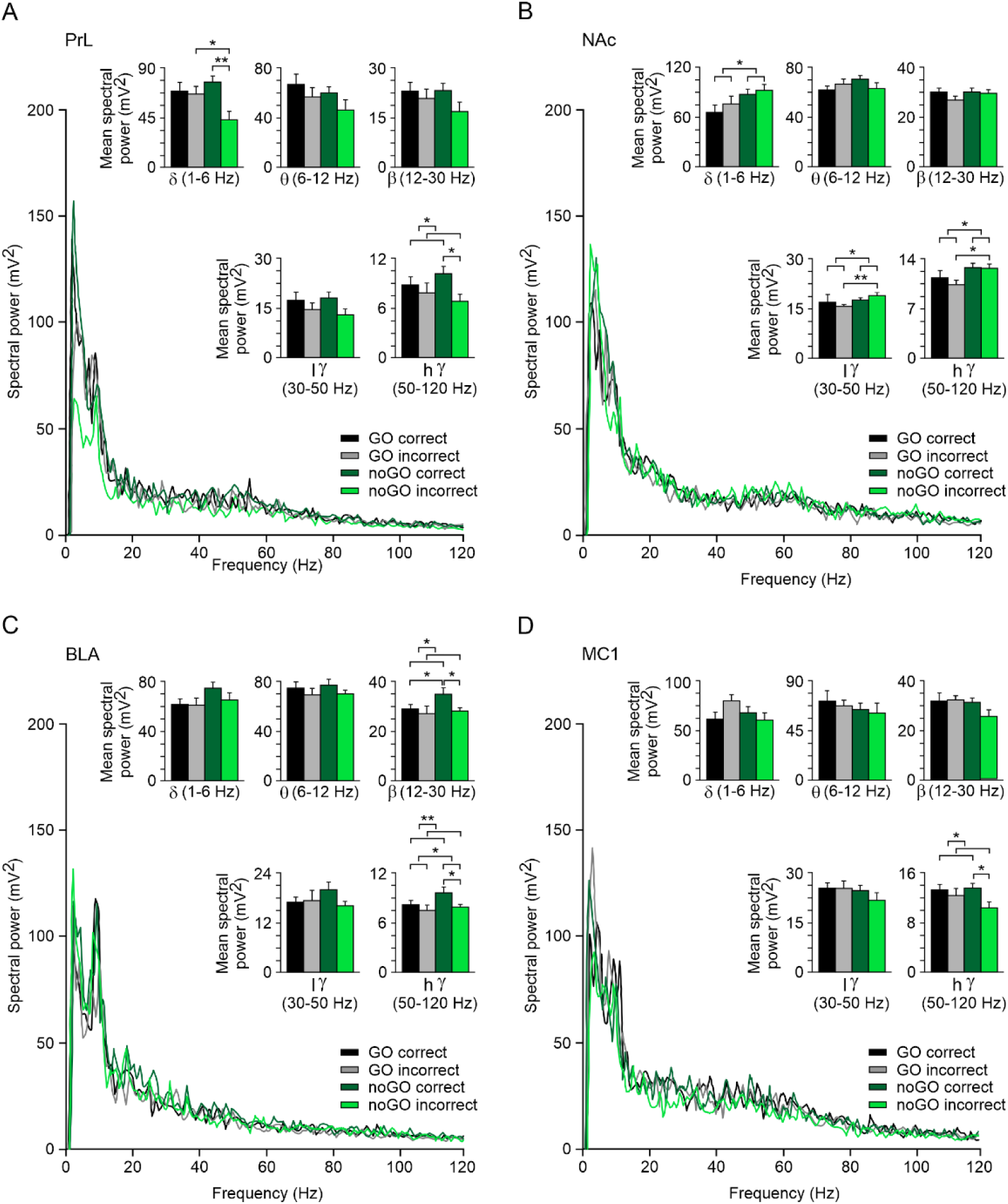
Spectral analyses of LFPs recorded in the PrL cortex, NAc, BLA, and MC1 during the performance of correct (black) and incorrect (gray) GO responses and correct (dark green) and incorrect (light green) noGO responses. **A-D.** LFP traces (n = 20) were collected from ≥ 5 animals. For GO correct responses, the selected traces were collected from the 2 s preceding touching the illuminated horizontal white rectangle, whereas for GO incorrect responses, traces were collected from the 2 s preceding the visual stimulus switching off. For noGO correct responses, traces were collected from the 2 s preceding the end of the visual stimulus (vertical green rectangle), whereas for incorrect responses, the selected traces were collected from the 2 s preceding a touch on the visual stimulus. Power spectra (1-120 Hz) corresponding to these four experimental situations are illustrated. Insets illustrate the power histograms for the five (delta, theta, beta, and low- and high-gamma) selected bands. Significant differences are indicated for the comparison of the four possible behaviors. Interestingly, significant differences were also observed when comparing correct (GO and noGO) vs. incorrect (GO and noGO) responses and GO (correct and incorrect) vs. noGO (correct and incorrect) responses. *, *p* < 0.05; **, *p* < 0.01.

**Figure 6.**
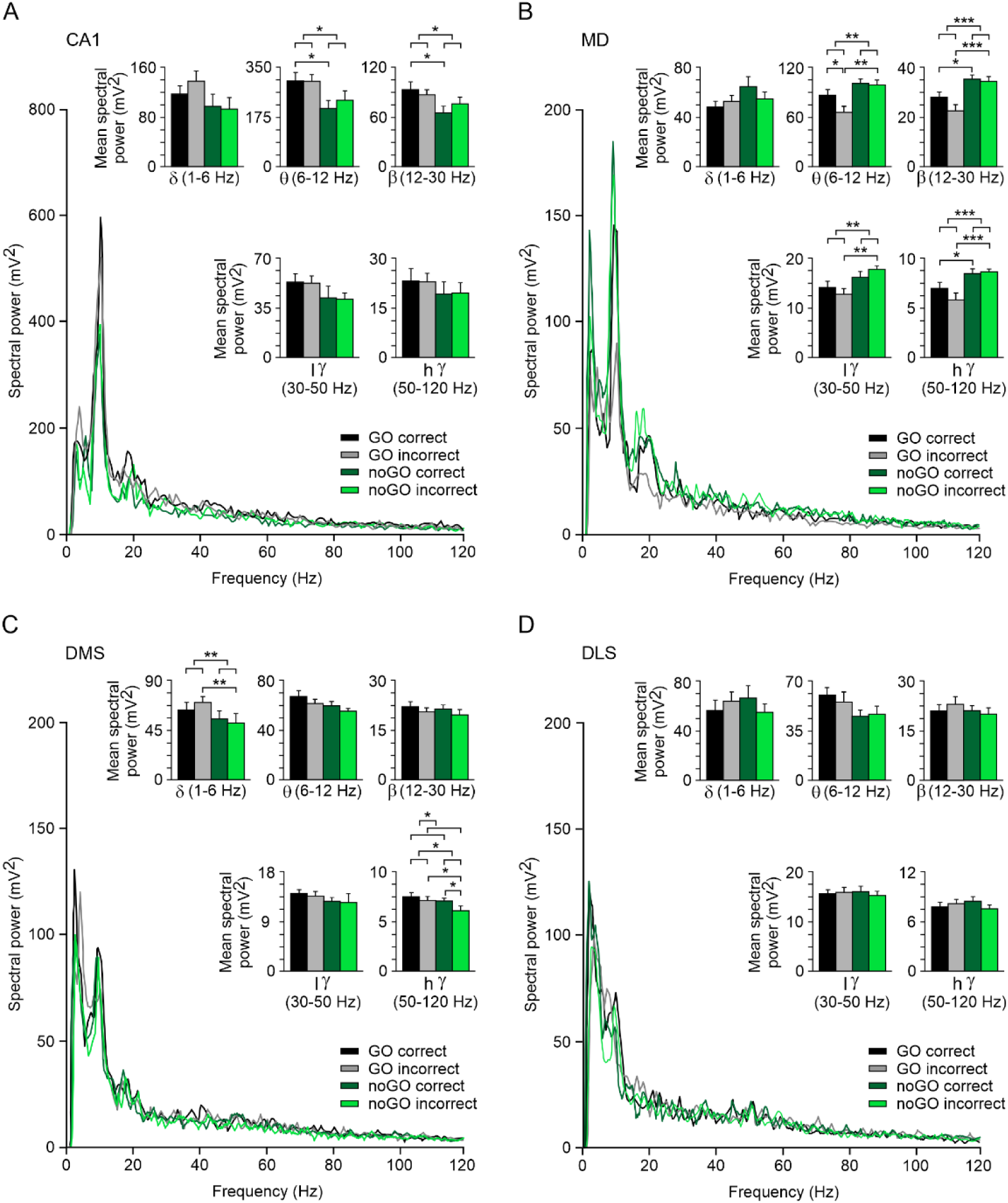
Spectral analyses of LFPs recorded in CA1, MD, DMS, and DLS during the performance of correct (black) and incorrect (gray) GO responses and correct (dark green) and incorrect (light green) noGO responses. **A-D**. LFP traces (n = 20) were collected from ≥ 5 animals. For GO correct responses the selected traces were collected from the 2 s preceding touching the illuminated white rectangle, whereas for GO incorrect responses, traces were collected from the 2 s preceding the visual stimulus being switched off. For noGO correct responses, traces were collected from the 2 s preceding the end of the visual stimulus (vertical green rectangle), whereas for incorrect responses, the selected traces were collected from the 2 s preceding touching the visual stimulus. Power spectra corresponding to the four experimental situations are illustrated. Insets illustrate the power histograms for the five (delta, theta, beta, and low- and high-gamma) selected bands. Significant differences are indicated for the comparison of the four behaviors. Interestingly, significant differences were also observed when comparing correct (GO and noGO) vs. incorrect (GO and noGO) responses and GO (correct and incorrect) vs. noGO (correct and incorrect). *, *p* < 0.05; **, *p* < 0.01; ***, *p* < 0.001.

In LFPs recorded in the PrL cortex (Fig. 5A), we observed that the spectral powers of the five selected bands were larger for correct GO and noGO responses than for incorrect ones, but the comparison between noGO correct vs. noGO incorrect responses was significantly different only for delta [*F*_(1,36)_ = 5.300, *p* = 0.027; *t* = 3.027; *p* < 0.005] and high-gamma bands [*F*_(1,36)_ = 4.670, *p* = 0.037; *t* = 2.388; *p* = 0.022]. Interestingly, the comparison of GO incorrect vs. noGO incorrect responses also yielded significant differences for the delta band [*F*_(1,36)_ = 5.300, *p* = 0.027; *t* = 2.072; *p* < 0.05], and comparing the two correct responses (GO and noGO) vs. the two incorrect ones yielded significant differences [*F*_(1,36)_ = 4.670, *p* = 0.037; *t* = 2.161; *p* = 0.037] for the high-gamma band.

In contrast, NAc recordings presented significantly larger powers for incorrect noGO vs. GO responses in the low-gamma [*F*_(1,36)_ = 3.354, *p* = 0.075; *t* = 3.174; *p* < 0.003] and high-gamma [*F*_(1,36)_ = 6.965, *p* = 0.012; *t* = 2.322; *p* = 0.026] bands. Also in this case, we observed significantly larger responses for correct and incorrect noGO responses vs. GO ones for delta [*F*_(1,36)_ = 4.899, *p* = 0.033; *t* = 2.213; *p* = 0.033], low-gamma [*F*_(1,36)_ = 7.062, *p* = 0.012; *t* = 2.657; *p* = 0.012], and high-gamma [*F*_(1,36)_ = 6.965, *p* = 0.012; *t* = 2.639; *p* = 0.012] bands (Fig. 5B).

As already described for LFPs recorded in the PrL cortex, LFPs corresponding to the BLA (Fig. 5C) presented larger power spectra for correct than for incorrect responses for the five selected bands, but were significantly different only for correct vs. incorrect noGO responses in the beta [*F*_(1,36)_ = 4.762, *p* = 0.036; *t* = 2.396; *p* = 0.022] and high-gamma [*F*_(1,36)_ = 7.539, *p* = 0.009; *t* = 2.353; *p* = 0.024] bands. The comparison of GO and noGO correct vs. incorrect responses also yielded significant differences in the beta [*F*_(1,36)_ = 4.762, *p* = 0.036; *t* = 2.182; *p* = 0.036] and high-gamma [*F*_(1,36)_ = 7.539, *p* = 0.009; *t* = 2.746; *p* = 0.009] bands. Finally, noGO (correct and incorrect) responses presented higher power than GO ones for the high-gamma band [*F*_(1,36)_ = 5.079, *p* = 0.030; *t* = 2.254; *p* = 0.030].

With respect to the MC1 (Fig. 5D), power spectra of the four experimental situations did not present a definite pattern, although correct responses presented larger powers than incorrect ones in the five selected bands, except for GO correct vs. incorrect responses in the delta band. Significant differences were found only between noGO correct vs. incorrect responses in the high-gamma band [*F*_(1,36)_ = 4.516, *p* = 0.041; *t* = 2.336; *p* = 0.025]. In addition, high-gamma power for correct GO and noGO responses was significantly larger than for incorrect responses [*F*_(1,36)_ = 4.516, *p* = 0.041; *t* = 2.125; *p* = 0.041].

LFPs recorded in the hippocampal CA1 area (Fig. 6A) presented larger power spectra for GO than for noGO correct responses for the five selected spectral bands, but were significantly different only for the theta [*F*_(1,36)_ = 7.190, *p* = 0.011; *t* = 2.329; *p* = 0.026] and beta [*F*_(1,36)_ = 5.797, *p* = 0.021; *t* = 2.452; *p* = 0.019] bands. Also, GO correct and incorrect responses taken together presented larger power spectra for theta [*F*_(1,36)_ = 7.190, *p* = 0.011; *t* = 2.681; *p* = 0.011] and beta [*F*_(1,36)_ = 5.797, *p* = 0.021; *t* = 2.408; *p* = 0.021] bands than the corresponding values for noGO correct and incorrect responses.

LFPs recorded in the MD presented a larger number of significant changes in the five selected spectral bands than any other brain site considered in the present study. The power spectra of the five bands were larger for noGO than for GO correct responses, reaching significant values for the beta [*F*_(1,36)_ = 20.348, *p* < 0.001; *t* = 2.444; *p* = 0.020] and high-gamma bands [*F*_(1,36)_ = 17.651, *p* < 0.001; *t* = 2.146; *p* = 0.039]. Also we noted that noGO correct and incorrect responses taken together reached larger spectral powers than the corresponding values for GO correct and incorrect responses; in this case, significant differences were found for theta [*F*_(1,36)_ = 12.386, *p* < 0.001; *t* = 3.519; *p* < 0.001], beta [*F*_(1,36)_ = 20.348, *p* < 0.001; *t* = 4.511; *p* < 0.001], low-gamma [*F*_(1,36)_ = 10.585, *p* = 0.002; *t* = 3.254; *p* = 0.002], and high-gamma [*F*_(1,36)_ = 17.651, *p* < 0.001; *t* = 4.201; *p* < 0.001] bands.

LFPs recorded in the DMS presented larger spectral powers for GO correct and incorrect responses than for the corresponding values of noGO correct and incorrect responses. These values were significantly different for delta [*F*_(1,36)_ = 9.238, *p* = 0.004; *t* = 3.039; *p* < 0.004] and high-gamma [*F*_(1,36)_ = 5.090, *p* = 0.030; *t* = 2.256; *p* = 0.030] bands. Other significant differences were found between GO incorrect and noGO incorrect responses also in the delta [*F*_(1,36)_ = 9.238, *p* = 0.004; *t* = 3.321; *p* < 0.002] and high-gamma [*F*_(1,36)_ = 5.090, *p* = 0.030; *t* = 2.376; *p* = 0.023] bands. In addition, high-gamma oscillations recorded during noGO correct responses presented a larger spectral power than the corresponding values for noGO incorrect responses [*F*_(1,36)_ = 5.706, *p* = 0.022; *t* = 2.469; *p* = 0.018]. Finally, high-gamma oscillations recorded during correct (GO and noGO) responses presented a larger spectral power than during incorrect ones [*F*_(1,36)_ = 5.706, *p* = 0.022; *t* = 2.389; *p* = 0.022].

LFPs recorded in the DLS did not present significant changes in the power of the five selected spectral bands during the performance of GO and noGO correct and incorrect responses.

Taken as a whole, present results indicate the presence of significant differences in the spectral power of LFPs recorded during the correct or incorrect performance of GO and noGO responses, corresponding mainly to the delta and gamma bands for some of the selected brain areas (PrL, NAc, and DMS) or including theta and/or beta and gamma bands for other structures (BLA, CA1, and MD). LFPs recorded in MC1 and DLS were not significantly modified in these experimental situations.

### The fundamental spectral components of LFP data were modified during the performance of GO and noGO correct and incorrect responses

In a following step, we performed a PSD analysis of LFPs collected during the performance of GO and noGO correct and incorrect responses. PSD analysis enabled determining the dominant periodicity of LFP recordings by quantifying the power peak at the frequency corresponding to that dominant periodicity (Jurado-Parras et al., 2013).

As illustrated in Fig. 7A,B, the dominant peaks of the PSD quantified from the 8 recording sites ranged within the values of the theta band (6–12 Hz): from 7.35 Hz computed for LFPs recorded in the DLS during GO incorrect responses to 10.15 Hz computed for LFPs recorded in CA1 during noGO incorrect responses. Four brain sites (BLA, CA1, MD and DMS) presented more-dominant peaks than the other four (PrL, NAc, MC1, and DLS).

**Figure 7.**
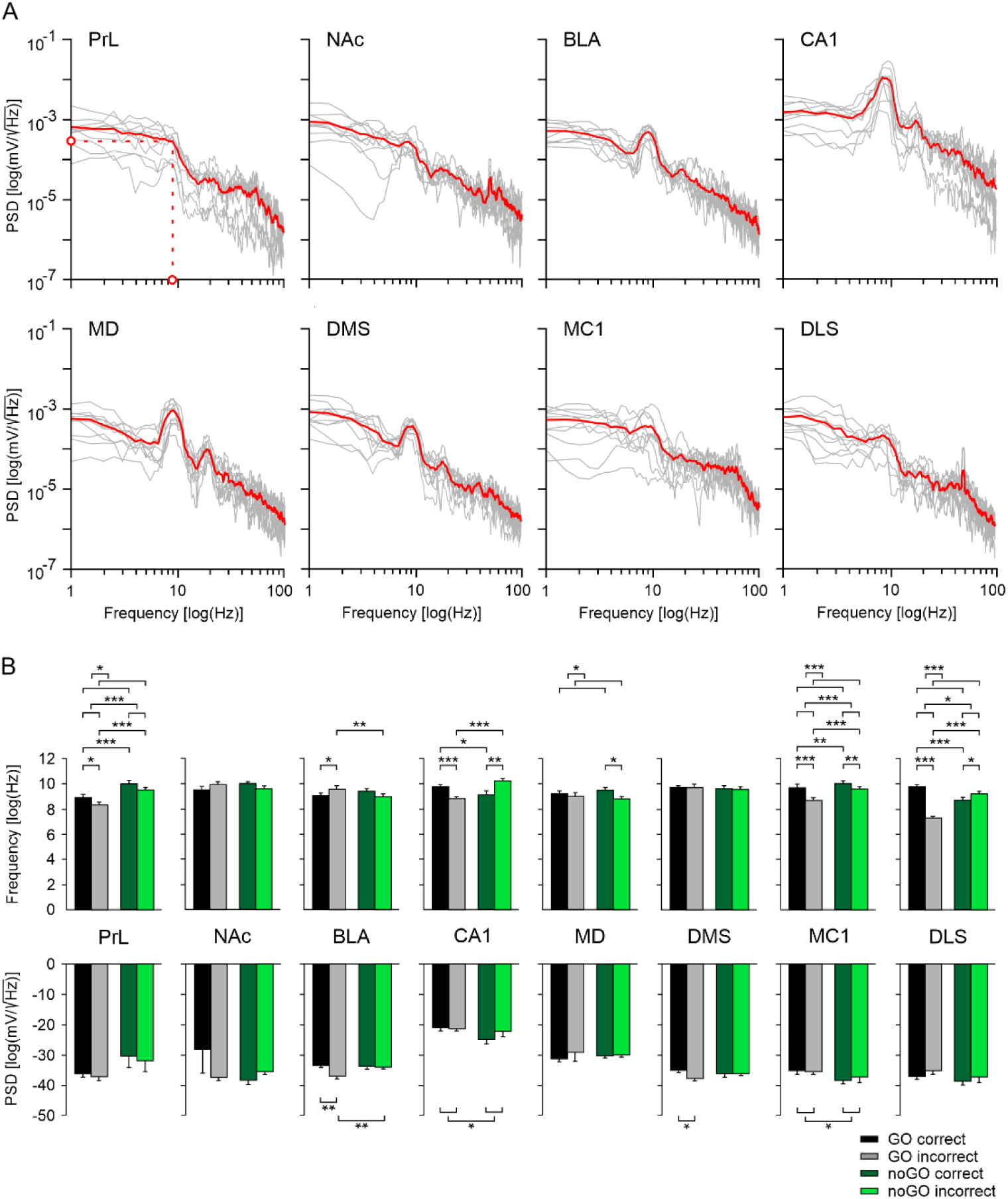
Differences in PSD for LFPs recorded in the 8 selected areas during the performance of correct and incorrect GO and noGO responses. **A**. PSD plots for LFP activities during the performance of GO correct responses. LFP traces (n = 20, from ≥ 5 animals) were collected from the 2 s preceding touching the illuminated white rectangle. Red traces represent mean values of collected traces. The red dotted lines indicate the fundamental components of the illustrated spectra (PSD peaks on the y-axis and their corresponding frequency on the x-axis). **B.** The top set of histograms represent mean frequency values for the fundamental spectral powers for the eight brain areas included in this study during the four experimental situations. Although all averaged PSDs presented peak values in the theta band, there were significant differences with the corresponding frequency values. The bottom set of histograms represent PSD values corresponding to the peak frequencies. Significant differences are indicated for the comparison of the four measured parameters between them. Interestingly, significant differences were also observed when comparing correct (GO and noGO) vs. incorrect (GO and noGO) responses and GO (correct and incorrect) vs. noGO (correct and incorrect). *, *p* < 0.05; **, *p* < 0.01; ***, *p* < 0.001.

LFPs recorded in the PrL presented significant differences in dominant frequencies between GO correct vs. incorrect responses [*F*_(1,36)_ = 8.084, *p* = 0.007; *t* = 2.489; *p* = 0.018], GO correct vs. noGO correct [*F*_(1,36)_ = 37.118, *p* < 0.001; *t* = 3.829; *p* < 0.001], and GO incorrect vs. noGO incorrect [*F*_(1,36)_ = 8.084, *p* = 0.007; *t* = 4.787, *p* < 0.001]. Additional significantly lower mean values were found between GO correct and incorrect vs. noGO correct and incorrect responses [*F*_(1,36)_ = 37.118, *p* < 0.001; *t* = 6.092, *p* < 0.001] and higher mean values for GO and noGO correct than for GO and noGO incorrect responses [*F*_(1,36)_ = 0.458, *p* = 0.503; *t* = 2.843, *p* = 0.007].

Dominant peak frequency quantified from LFPs recorded in the hippocampal CA1 area was significantly larger for GO correct compared with GO incorrect responses [*F*_(1,36)_ = 25.599, *p* < 0.001; *t* = 3.665, *p* < 0.001], noGO incorrect vs. correct responses [*F*_(1,36)_ = 25.599, *p* < 0.001; *t* = 3.490, *p* < 0.001], GO correct vs. noGO correct responses [*F*_(1,36)_ = 25.599, *p* < 0.001; *t* = 2.269, *p* = 0.029], and noGO incorrect responses vs. GO incorrect responses [*F*_(1,36)_ = 25.599, *p* < 0.001; *t* = 4.887, *p* < 0.001].

LFPs recorded in the MC1 presented numerous changes in the PSD dominant peak frequency. Significantly larger frequencies were observed for GO correct vs. incorrect responses [*F*_(1,36)_ = 27.771, *p* < 0.001; *t* = 4.046, *p* < 0.001], noGO correct vs. incorrect responses [*F*_(1,36)_ = 27.771, *p* < 0.001; *t* = 3.407, *p* = 0.002], noGO correct vs. GO correct responses [*F*_(1,36)_ = 24.688, *p* < 0.001; *t* = 3.194, *p* = 0.003], noGO incorrect vs. GO incorrect responses [*F*_(1,36)_ = 24.688, *p* < 0.001; *t* = 3.833, *p* < 0.001], noGO correct and incorrect vs. GO correct and incorrect responses [*F*_(1,36)_ = 24.688, *p* < 0.001; *t* = 4.969, *p* < 0.001], and GO and noGO correct vs. GO and noGO incorrect responses [*F*_(1,36)_ = 27.771, *p* < 0.001; *t* = 5.270, *p* < 0.001].

The LFPs recorded in the DLS also presented significant changes of the dominant peak frequency for many comparisons between the different variables considered in this study. Significantly larger values were observed between GO correct vs. incorrect responses [*F*_(1,36)_ = 27.319, *p* < 0.001; *t* = 9.519, *p* < 0.001], noGO incorrect vs. correct responses [*F*_(1,36)_ = 27.319, *p* < 0.001; *t* = 2.198, *p* = 0.034], GO correct vs. noGO correct responses [*F*_(1,36)_ = 69.466, *p* < 0.001; *t* = 4.395, *p* < 0.001], noGO incorrect vs. GO incorrect responses [*F*_(1,36)_ = 69.466, *p* < 0.001; *t* = 7.392, *p* < 0.001], GO correct and incorrect vs. noGO correct and incorrect responses [*F*_(1,36)_ = 4.490, *p* = 0.041; *t* = 2.119, *p* = 0.041], and GO and noGO correct vs. GO and noGO incorrect responses [*F*_(1,36)_ = 4.490, *p* = 0.041; *t* = 5.227, *p* < 0.001].

Two of the selected brain sites (NAc and DMS) did not present significant changes in frequency or power of their dominant peak frequencies. BLA presented significantly larger values in frequency only for GO incorrect vs. GO correct responses [*F*_(1,36)_ = 8.762, *p* = 0.005; *t* = 2.193, *p* = 0.035] and for GO incorrect and noGO incorrect responses [*F*_(1,36)_ = 8.762, *p* = 0.005; *t* = 3.189, *p* = 0.003]. Similarly, the MD presented significantly larger dominant frequencies only for noGO correct vs. incorrect responses [*F*_(1,36)_ = 4.257, *p* = 0.046; *t* = 2.403, *p* = 0.022].

### Spectrograms of the selected LFP segments during GO and noGO stimulus presentations illustrate the different roles played by the selected brain sites in this instrumental learning task

The spectral analysis of the selected 2-second LFP segments illustrated in Figs. 5-7 yielded interesting details of the different contributions of the 8 brain areas during the correct or incorrect performance of GO and noGO tasks. But for a more detailed understanding of the sequential changes in spectral power across the indicated time it is also useful to take advantage of time-frequency representations. In Fig. 8 are illustrated the spectrograms averaged from 20 samples of 2-second segments collected from ≥ 5 rats from LFPs recorded in the PrL cortex. Illustrations correspond to GO correct (Fig. 8A) and incorrect (Fig. 8B) responses and to noGO correct (Fig. 8C) and incorrect (Fig. 8D) responses. Illustrated spectrograms correspond to the 2 s preceding GO correct and noGO incorrect responses (Fig. 8A,D) or to the switching off of the GO incorrect and noGO correct stimuli (Fig. 8B,C).

**Figure 8.**
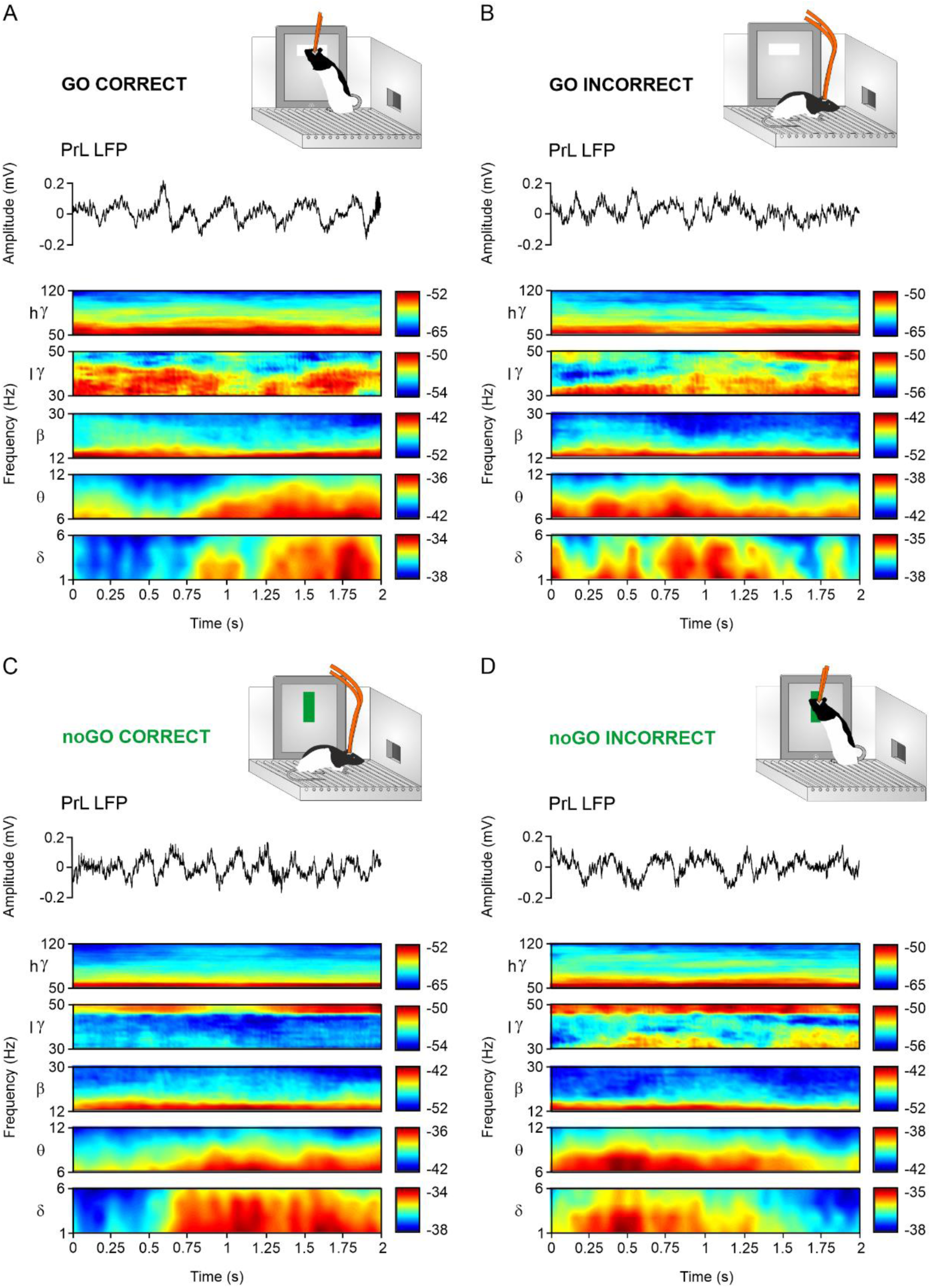
Dynamic changes in LFP oscillations recorded in the PrL cortex during the four experimental situations. **A-D**. At the top are illustrated the typical animal behaviors observed during GO correct (**A**) and incorrect (**B**), and noGO correct (**C**) and incorrect (**D**) responses. Below are illustrated a representative LFP trace and multitaper time-frequency representations averaged from 20 traces (2-second segment each) collected from ≥ 5 animals. Note the similarity of delta and theta bands for GO and noGO correct responses vs. those presented during GO and noGO incorrect responses. The frequency range of the selected bands is indicated to the left and the respective color calibration bar is indicated to the right of the corresponding spectrogram.

The illustrated spectrograms present a dominant spectral power corresponding to delta and theta bands occurring very closely to the correct responses for GO and noGO correct responses (Fig. 8A,C) and the opposite phenomenon for the incorrect ones (Fig. 8B,D). These results suggest that these increases in power were related more to the appropriate selection of the correct response than to the programming of a motor behavior to approach the iPad and to touch the displayed visual stimulus. Fig. 9 illustrates a similar set of spectrograms collected from the other 7 brain regions included in this study. Contrary to what we observed in PrL spectrograms, both CA1 and MC1 presented higher spectral powers for delta and theta bands preferentially related to active behaviors to approach the visual stimulus during GO correct and noGO incorrect responses and a decrease in power for both bands during GO incorrect and noGO correct responses. Although not as well defined, spectrograms collected from DLS presented profiles similar to those already mentioned for CA1 and MC1. Finally, spectrograms collected from the rest of the implanted sites (NAc, BLA, MD, and DMS) presented similar power values across the 2-second epochs and for the four experimental situations.

**Figure 9.**
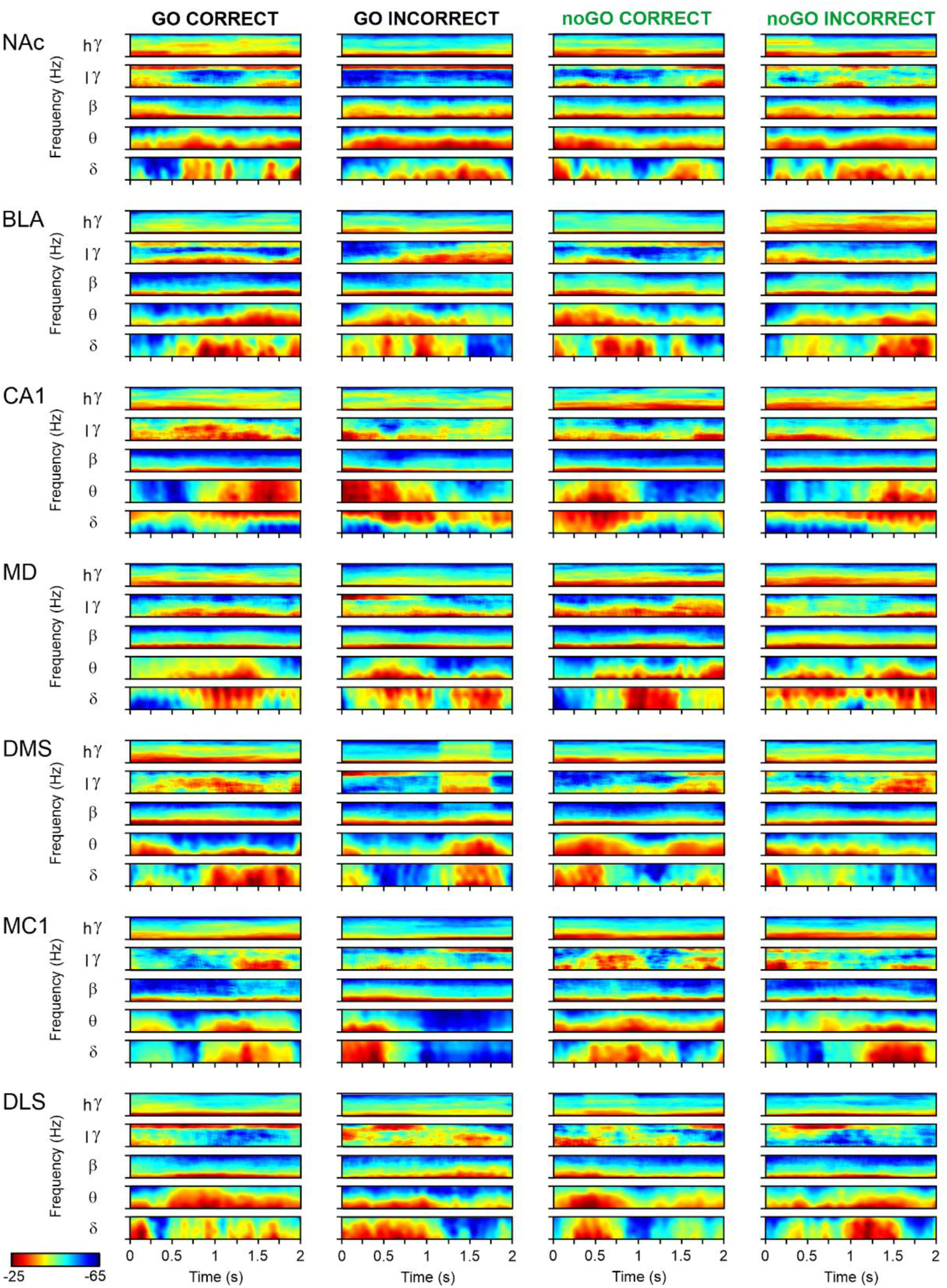
Dynamic changes in LFP oscillations recorded in 7 selected brain areas during the four experimental situations. From top to bottom are illustrated the spectrogram collected from LFPs recorded in the indicated areas (NAc, BLA, CA1, MD, DMS, MC1, and DLS) during the four experimental situations (GO correct, GO incorrect, noGO correct, and noGO incorrect). As indicated in Fig. 8, multitaper time-frequency representations were averaged from 20 traces collected from ≥ 5 animals. Note the different power of five selected frequency bands (see their ranges in Fig. 8) during the performance of the four experimental situations. A common color calibration bar is illustrated at the bottom left.

### Analysis of coherence between the PrL cortex and the other seven recorded sites for the four experimental situations

An interesting question was to determine how the PrL cortex was functionally interconnected with the other recording sites during the moment the experimental rats decided whether or not to touch the iPad screen as a correct (or incorrect) response to the displayed GO or noGO visual stimuli. For two representative relationships, in Fig. 10A are illustrated coherences presented between PrL and NAc (Fig. 10A, left two displays) and PrL and BLA (Fig. 10A, right two displays) for two of the experimental situations considered in the present study (GO and noGO correct responses) during representative trials. As illustrated in Fig. 10A, a variable coherence (0-0.5) was observed for these two relationships (PrL-NAc and PrL-BLA) during the 2 s preceding the definite decision made by the experimental animal. In Fig. 10B,C are illustrated mean values for the coherence coefficient determined from the average of 20 LFP segments, selected from ≥ 5 animals, during the 2 s preceding the final decision taken by the experimental animals with respect to the stimuli presented on the iPad screen. In this case, the four experimental situations are represented. There were many different moments during which coherence between the selected 7 pairs of recordings (PrL-NAc, PrL-BLA, PrL-CA1, PrL-MD, PrL-DMS, PrL-MC1, and PrL-DLS) was modified. Coherence values ≥ 0.4 (Fig. 10B, dotted red lines) were found between PrL-NAc (delta and theta bands), PrL-CA1 (theta band), PrL-DMS (delta band), PrL-MC1 (theta and low-gamma bands), and PrL-DLS (delta and theta bands). In addition, there was an experimental situation (noGO correct responses) in which coherence analysis for the theta band reached values > 0.4 for different pairs of brain sites (PrL-NAc, PrL-CA1, PrL-MC1, and PrL-DLS) (Fig. 10C). These results indicate a high functional adherence between the PrL region and the other brain sites included in this study, during the inhibition of a motor behavior in order to correctly perform the task.

**Figure 10.**
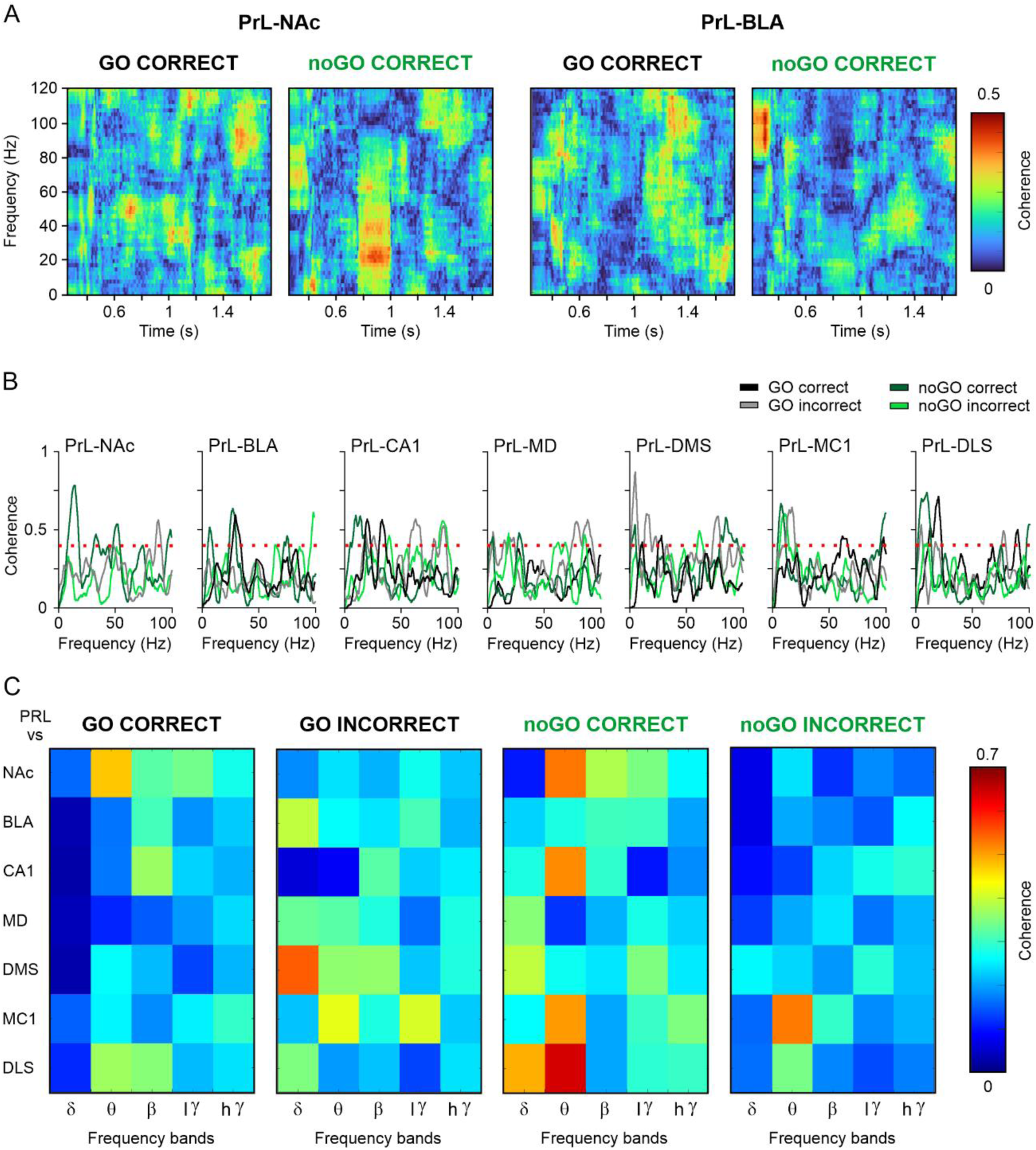
Spectrograms representing the coherence between the PrL cortex and the other recording sites included in this study. **A**. Representative maps of the time-frequency coherence between PrL and NAc (left set of spectrograms) and PrL and BLA (right set of spectrograms). Illustrated maps correspond to 20 samples (2-second segment each) collected from ≥ 5 rats. Note that high coherence between compared spectrograms appeared at different moments and frequencies. **B**. Coherence profiles for the four experimental situations (GO correct and incorrect and noGO correct and incorrect) between PrL spectrograms and those collected from the other seven recording sites. Note the presence of high coherence values (> 0.4) between PrL and the other recording sites in different frequency bands (dotted red lines). **C**. A comparative analysis of coherence between PrL cortex and the other recording sites for the four experimental situations. Note the high coherence between PrL and other (NAc, CA1, MC1, and DLS) recorded areas during noGO correct responses.

### Changes in spectral power for four selected brain sites comparing LFPs recorded in the initial vs. the final training sessions

Across the successive conditioning sessions, we noted that the spontaneous LFP activities were different comparing the beginning with the end of the session, and also between LFPs recorded from initial sessions to those collected in later ones. In order to carry out a quantitative analysis of this observation, we selected 20-second segments from the initial and final part of the 1st and 2nd conditioning sessions [i.e., before any trace of associative learning (Fig. 3A)] vs. the 14th and 18th sessions [i.e., when the animals have already acquired the GO, noGO task] (Fig. 3E,G)]. In Fig. 11 are illustrated the experimental design (Fig. 11A) and representative examples of LFP segments recorded in the PrL from the initial and final part of sessions 1 and 18 from the same subjects. The increase in delta/theta oscillations during the final part of the 18th session can be easily observed (Fig. 11B). We carried out a spectral analysis of traces collected from two recorded areas (PrL and NAc) preferentially related with cognitive aspects of the learning task (Fig. 11C,D) and another two areas (DLS and CA1) more related with the sensorimotor activities involved in the elaboration of the corresponding motor responses (Fig. 11E,F). The aim was to look for differences in the spectral power of LFP oscillations collected from these four brain sites.

**Figure 11.**
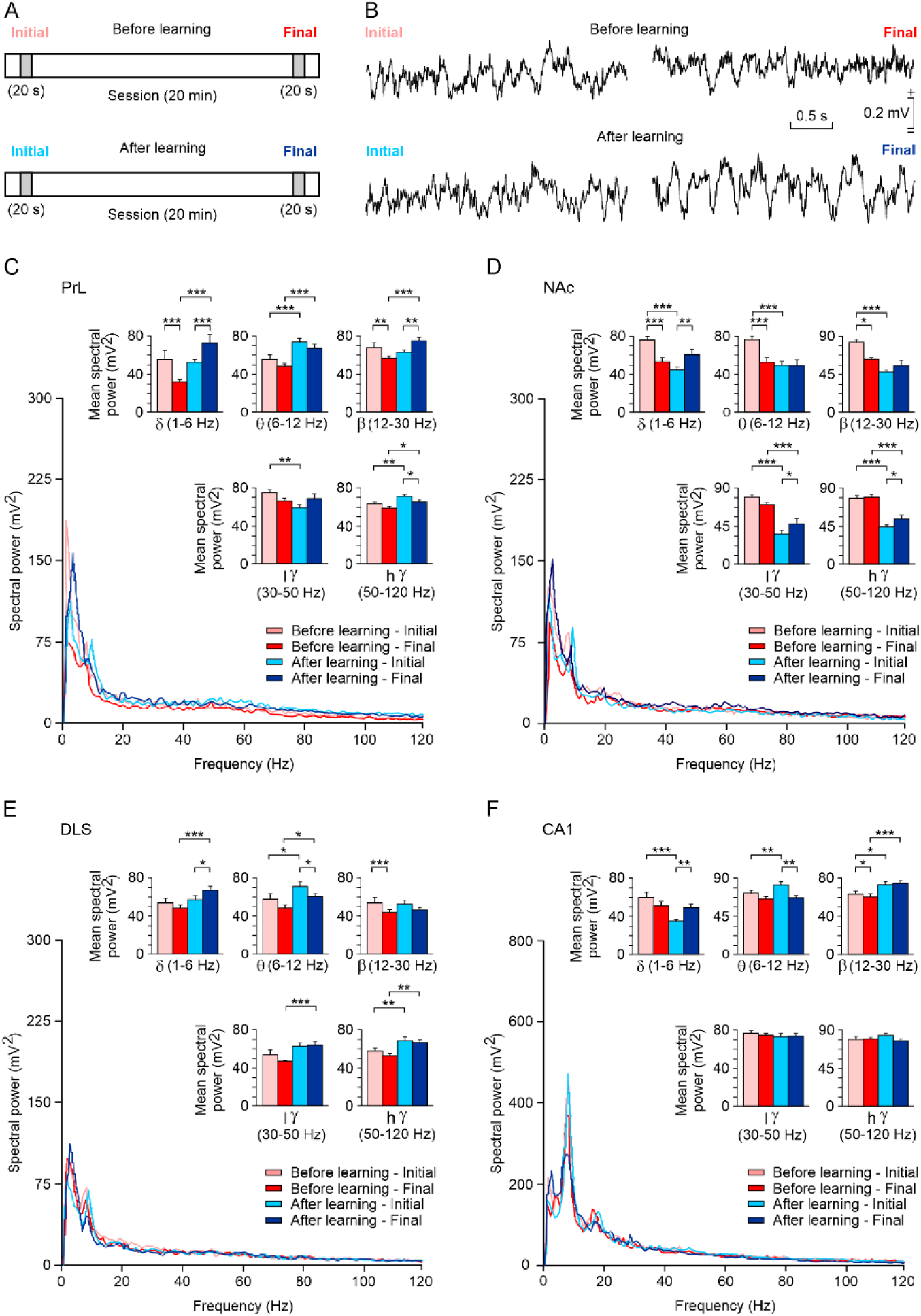
A comparison of the spectral power of LFPs collected from the beginning and the end of training sessions that took place before and after the acquisition of the GO/noGO task. **A**. We collected a total of eight 20-second samples of LFPs recorded before (sessions 1 and 2) and after (sessions 14 and 18) proper acquisition of the GO/noGO task. The two latter sessions included GO and noGO presentations. For comparative purposes, selected LFP traces were collected from the initial and the final parts of the recording sessions. **B**. Representative traces collected from the initial and final parts of the recording sessions from the PrL cortex, before and after learning. Note the presence of noticeable low-frequency (delta/theta) waves in the LFP trace collected from the final part of a recording session carried out after learning. **C-F**. Spectral analyses of LFPs recorded before (initial and final) and after (initial and final) the acquisition of the GO/noGO task from PrL, NAc, DLS, and CA1 recording sites. Note the significant changes in spontaneous LFPs taking place at the different recording sites before and after learning and in the initial and final part of recording sessions. *, *p* < 0.05; **, *p* < 0.01; ***, *p* < 0.001.

LFPs recorded in the PrL cortex presented evident differences for traces collected from the first and final parts of the same session and comparing sessions before and after the acquisition of the learning task. In general, LFP oscillations presented a decrease in power from the initial to final segments during before-learning sessions, and the opposite during after-ones. For the delta band, we noted significantly larger values between initial vs. final parts during the before-learning sessions [*F*_(1,19)_ = 57.052, *p* < 0.001; *t* = 5.133, *p* < 0.001], and the opposite for after-learning ones [*F*_(1,19)_ = 57.052, *p* < 0.001; *t* = 4.981, *p* < 0.001]. Also, we observed the presence of significantly lower values comparing the spectral powers of segments collected from the final part of sessions recorded before vs. after learning [*F*_(1,19)_ = 57.052, *p* < 0.001; *t* = 8.478, *p* < 0.001]. For theta bands, we found significantly lower spectral powers for the two initial [*F*_(1,19)_ = 25.214, *p* < 0.001; *t* = 3.477, *p* < 0.001] and the two final [*F*_(1,19)_ = 25.214, *p* < 0.001; *t* = 3.785, *p* < 0.001] segments. Beta bands presented significantly larger values for initial vs. final parts during before-learning sessions [*F*_(1,19)_ = 23.516, *p* < 0.001; *t* = 3.367, *p* = 0.002], and the opposite for after-learning ones [*F*_(1,19)_ = 23.516, *p* < 0.001; *t* = 2.854, *p* = 0.0017]. Also, we found the presence of significant differences comparing the spectral powers of segments collected from the final part of sessions recorded before vs. after learning [*F*_(1,19)_ = 4.049, *p* = 0.059; *t* = 4.420, *p* < 0.001]. The low-gamma band presented significantly larger values for the initial segments recorded before vs. after learning [*F*_(1,19)_ = 4.967, *p* = 0.038; *t* = 2.229, *p* = 0.038]. Finally, high-gamma bands presented significantly larger values for initial vs. final recordings during after-learning sessions [*F*_(1,19)_ = 26.709, *p* < 0.001; *t* = 2.204, *p* = 0.034], lower values for initial segments from before-learning vs. after-learning sessions [*F*_(1,19)_ = 26.709, *p* < 0.001; *t* = 3.276, *p* = 0.002], and also lower values for final segments from before-learning vs. after-learning sessions [*F*_(1,19)_ = 26.709, *p* < 0.001; *t* = 3.276, *p* = 0.002].

As already indicated for LFPs collected from the PrL cortex, LFP segments collected from the NAc presented a decrease in power from the initial to final segments during before-learning sessions and the opposite during after-learning ones. For the delta band, we observed significantly larger values for initial vs. final parts during before-learning sessions [*F*_(1,19)_ = 32.048, *p* < 0.001; *t* = 4.258, *p* < 0.001], and the opposite for after-learning ones [*F*_(1,19)_ = 32.048, *p* < 0.001; *t* = 3.198, *p* = 0.003]. Also, we noted the presence of significant differences comparing the spectral powers of segments collected from the initial part of sessions recorded before vs. after learning [*F*_(1,19)_ = 7.768, *p* = 0.012; *t* = 5.749, *p* < 0.001]. For the theta band, we collected larger spectral powers for the initial vs. final LFPs collected during before-learning sessions [*F*_(1,19)_ = 12.301, *p* = 0.002; *t* = 3.819, *p* < 0.001] and between initial recordings collected during the before-learning and after-learning periods [*F*_(1,19)_ = 11.743, *p* = 0.003; *t* = 4.841, *p* < 0.001]. For the beta band, we observed larger values for the initial vs. the final LFPs collected during before-learning sessions [*F*_(1,19)_ = 26.441, *p* < 0.001; *t* = 3.753, *p* < 0.001] and for the initial recordings collected during the before-learning and after-learning sessions [*F*_(1,19)_ = 32.901, *p* < 0.001; *t* = 7.625, *p* < 0.001]. The low-gamma band presented significantly lower values for initial vs. final LFP traces collected during the after-learning period [*F*_(1,19)_ = 8.772, *p* = 0.008; *t* = 2.219, *p* = 0.033], and larger values for initial LFP traces collected from before- vs. after-learning sessions [*F*_(1,19)_ = 81.315, *p* < 0.001; *t* = 8.552, *p* < 0.001], and during final LFP traces collected from before- and after-learning sessions [*F*_(1,19)_ = 81.315, *p* < 0.001; *t* = 4.446, *p* < 0.001]. Finally, the high-gamma band also presented slightly lower values for initial vs. final LFP traces collected during the after-learning period [*F*_(1,19)_ = 3.273, *p* = 0.086; *t* = 2.039, *p* = 0.049], and larger values for initial LFP traces collected from before- vs. after-learning sessions [*F*_(1,19)_ = 90.732, *p* < 0.001; *t* = 7.578, *p* < 0.001], and for final LFP traces collected from before- vs. after-learning sessions [*F*_(1,19)_ = 90.732, *p* < 0.001; *t* = 5.932, *p* < 0.001].

In general, LFPs recorded from the DLS presented larger spectral power for traces recorded after learning than before. For the delta band, we found significant differences between initial vs. final traces recorded during the after-learning period [*F*_(1,19)_ = 5.254, *p* = 0.033; *t* = 2.172, *p* = 0.036] and between the final traces recorded before- and after-learning sessions [*F*_(1,19)_ = 9.670, *p* = 0.006; *t* = 3.858, *p* < 0.001]. For the theta band, significantly larger values were found between initial and final traces recorded during the after-learning period [*F*_(1,19)_ = 9.693, *p* = 0.006; *t* = 2.375, *p* = 0.023], between the two initial traces from before- vs. after-learning periods [*F*_(1,19)_ = 10.606, *p* = 0.004; *t* = 2.515, *p* = 0.016], and between the final traces recorded before- and after-learning sessions [*F*_(1,19)_ = 10.606, *p* = 0.004; *t* = 2.119, *p* = 0.041]. The beta band presented significantly larger values only when comparing initial vs. final LFP traces collected before-learning [*F*_(1,19)_ = 12.821, *p* = 0.002; *t* = 3.483, *p* < 0.001]. The low-gamma band presented significantly lower values only between final traces recorded before- vs. after-learning [*F*_(1,19)_ = 15.262, *p* < 0.001; *t* = 3.520, *p* = 0.001]. Finally, the high-gamma band presented significantly lower values between initial LFP traces recorded from before- vs. after-learning sessions [*F*_(1,19)_ = 31.146, *p* < 0.001; *t* = 2,984, *p* = 0.005], and the opposite during final traces collected from before- vs. after-learning sessions [*F*_(1,19)_ = 31.146, *p* < 0.001; *t* = 3.188, *p* = 0.003].

Recordings collected from the CA1 area during training sessions carried out before and after the acquisition of the GO/noGO task presented significant differences for delta, theta, and beta bands. For the delta band, we observed a significant increase in power of final vs. initial traces collected from after-learning sessions [*F*_(1,19)_ = 14.442, *p* < 0.001; *t* = 2.733, *p* = 0.010] and a decrease when comparing the initial LFPs collected from before- and after-learning sessions [*F*_(1,19)_ = 17.665, *p* < 0.001; *t* = 5.662, *p* < 0.001]. For the theta band, we observed a significant decrease in power of final vs. initial traces collected from after-learning sessions [*F*_(1,19)_ = 7.905, *p* = 0.011; *t* = 3.325, *p* = 0.002] and an increase when comparing the initial LFPs collected from before- and after-learning sessions [*F*_(1,19)_ = 4.161, *p* = 0.056; *t* = 2.703, *p* < 0.010]. Finally, for the beta band, we observed a significant increase from initial LFPs comparing recordings carried out before- and after-learning [*F*_(1,19)_ = 27.501, *p* < 0.001; *t* = 2.333, *p* = 0.026] and between final recordings also comparing recordings carried out before- vs. after-learning [*F*_(1,19)_ = 27.501, *p* < 0.001; *t* = 3.616, *p* < 0.001].

## DISCUSSION

The present results convincingly show that the PrL participates actively in the proper election of GO and noGO responses and less definitely in the incorrect ones. This is an indication that the PrL cortex is primarily involved in cognitive and motivation aspects of this instrumental task and not in the execution of reward-directed behaviors. Other brain areas (CA1, MC1, DLS) were more involved with the performance of the necessary motor sequences to reach the iPad and touch the displayed visual stimulus. The other four areas (NAc, BLA, MD, and DMS) presented spectral powers with not so-defined profiles during the four types of possible response to stimulus presentations. Nevertheless, some structures, such as the NAc, were specifically related to the acquisition of the instrumental task, being related to the learning task in moments outside stimulus presentations. Below is presented a detailed description of the functional characteristics of the eight brain sites selected in this study during the performance of the learning task.

### Main contributions of the different brain areas selected in this study

The mPFC, and particularly, the PrL cortex, is clearly involved in goal-directed activities, decision-making tasks, and social interactions. As already reported and further confirmed here, LFP oscillations recorded in the PrL cortex modulate their delta/theta spectral powers depending on the required cognitive and/or behavioral activities. As shown here, and in accordance with previous studies (Capuzzo and Floresco, 2020), the PrL is also involved in the selection of the proper decisions for GO correct and noGO correct responses. As illustrated in Fig. 8, spectrograms of LFPs recorded during the four experimental situations support the proposal that PrL activities are more directly related to cognitive processes than to sensorimotor commands, although the proper selection of the most-appropriate motor activities are also under its indirect control (Leal-Campanario et al., 2007). Unitary recordings of mPFC neurons carried out in rats during GO/noGO tests further support these contentions (de Haan et al., 2018). In addition, Karalis et al. (2016) have reported the presence of 4-Hz oscillations in the dorsal mPFC during fear behaviors. Similar frequency oscillations (4-6 Hz) have been reported in rats when approaching a feeder to collect a food reward (Hernández-González et al., 2017) and during the cooperative acquisition of an instrumental learning task also in rats (Conde-Moro et al., 2019). This dominant delta band apparently helps to synchronize the activities between prefrontal, hippocampal, and ventral tegmental areas (Fujisawa and Buzsáki, 2011), a fact further confirmed here with respect to other brain areas for cognitive (NAc, BLA, and DMS) and motor/behavioral (MC1, MD, CA1, and DLS) activities. In this respect, it is generally assumed that oscillatory brain activities, in a wide range of bands, serve to coordinate neuronal firing in neural networks distributed across different cortical and subcortical areas (Buzsáki and Draguhn, 2004). For example, different frequencies of brain oscillations (from delta to high-gamma) are evoked during instrumental learning, social interactions, fearful situations, and other motor displays (Fujisawa and Buzsáki, 2011; Cebolla and Cheron, 2015; Tendler and Wagner, 2015; Karalis et al., 2016; Hernández-González et al., 2017).

Present results further confirm that the PrL is functionally related in diverse ways with the other areas included in this study. To start with, the NAc is highly interconnected with the mPFC and is particularly involved in the generation of reward-seeking behaviors (Nicola, 2010; Floresco, 2015). NAc lesions and/or experimental manipulations of its dopamine inputs decrease seeking behaviors in response to incentive stimuli during different types of associative learning task (Nicola, 2010; Corbit and Balleine, 2011; Fraser and Janak, 2017). It has been confirmed recently that core NAc neurons are preferentially related to incentive vs. instructive stimuli (Sicre et al., 2020). The present results convincingly show the involvement of the NAc in GO and (mainly) noGO correct responses.

The amygdala—particularly the BLA nucleus—has classically been considered a structure involved in the acquisition of conditioned fear responses and as a detector of actual or potential threats to survival (LeDoux, 2000). In addition, the amygdaloid complex seems to be relevant to the presence of emotional expressions and other types of noticeable stimulus. For example, in a GO/noGO test carried out in humans, bilateral amygdala activation was more evident for relevant GO and noGO stimuli (Ousdal et al, 2008). It has been shown recently that the acquisition of a cued fear conditioning in mice is dependent on the proper activity of bidirectional glutamatergic projections between mPFC and BLA circuits, and that presynaptic mPFC and postsynaptic BLA NMDARs are required for fear memory formation, although not for its expression (Bertocchi et al., 2023). Neuronal-astrocyte synergy has also been proposed to underlie long-term fear memory formation (Sun et al., 2024). Although only positive rewards were presented in this study, the BLA was preferentially active for noGO vs. GO correct responses, suggesting an important contribution of this brain center to the inhibitory component necessary for the generation of correct noGO responses.

Although the dorsal striatum seems to be involved in the acquisition and proper performance of operant learning tasks, there are important functional differences between striatal regions. Thus, the DMS is preferentially connected with the mPFC, and mostly involved in the associations between responses and outcomes and the motivational control of goal-directed actions. In contrast, the DLS receives sensorimotor information from the respective cortical areas, and is more related to the proper formation of skilled movements and motor habits by the appropriate association between stimuli and responses (Stalnaker et al., 2010; Graybiel and Grafton, 2015; Corbit and Janak, 2016; Lipton et al., 2019; Stubbendorff et al., 2019; Vandaele et al., 2019). Based on simultaneous unitary recordings from PrL and dorsal striatum neurons, Stubbendorff et al. (2019) propose that this functional network also underlies reward-related decisions. Present results suggest an important functional connectivity between the PrL cortex and the dorsal striatum. It has already been suggested that repetitive behavioral sequences continue to engage the unitary activities of both (DLS, DMS) striatal regions for repeated presentation of the same demanded associative learning task (Vandaele et al., 2019). Moreover, a shift in control from dorsomedial to dorsolateral striatum during skill and habit formation has been reported (Turner et al., 2022). Since LFPs were recorded during the final set of sessions when the expected outcomes were well established, it could be thought that the GO/noGO test presented particular attentional and behavioral difficulties. Nevertheless, LFPs recorded in the DMS were more modified in spectral power than those collected from the DLS (Fig. 6). In the same sense, coherence between PrL and DMS reached higher values (> 0.4) than those collected for PrL-DLS (Fig. 11).

Because of its extensive cortical and subcortical network of afferent and efferent pathways, the motor cortex represents the final site for acquired motor commands ready to be transferred to brainstem and spinal motoneurons (Grillner and El Manira, 2020). However, it has already been shown in mice (Hasan et al., 2013) and rabbits (Ammann et al., 2016) that motor cortex circuits actively participate in the acquisition of both classical and instrumental conditioning tasks. In addition, mPFC projections play an inhibitory control on motor cortex, and related, circuits. Thus, electrical stimulation of the mPFC prevents the expression of newly acquired conditioned eyelid responses (Leal-Campanario et al., 2007). Although the activity of the MC1 was not specifically related to GO/noGO responses (Fig. 9), there were evident relationships between PrL and MC1 in the changes in the dominant peak frequencies collected for the four experimental situations considered in the present study (Fig. 7).

Prefrontal connections with the thalamic MD nucleus constitute a useful criterion for the proper identification of prefrontal areas in different species of mammals (Fuster, 2001; Leal-Campanario et al., 2007). MD functions have been related to attentional states, behavioral planning, and the proper maintenance of ongoing working memories (Bolkan et al., 2017). The MD was significantly more active during noGO vs. GO correct and incorrect responses for theta, beta, and low- and high-gamma bands, suggesting a particular contribution to the special attentional requirements for the generation of noGO responses.

The hippocampus has been reported to be involved in the proper representation of spatial coordinates and spatiotemporal sequences (Moser and Moser, 2013; Buzsáki and Llinás, 2017), but also in declarative memories and in some types of associative learning task, such as the classical conditioning of nictitating membrane/eyelid responses (Thompson, 1988; Gruart et al., 2006). In contrast, and apart from a specific relationship with the expression of consummatory behaviors, the hippocampus does not seem to be particularly involved in the acquisition of instrumental conditioning tasks (Jurado-Parras et al., 2013). Present results indicated a decline in theta and beta powers for noGO tasks (Fig. 6) that cannot be ascribed to the accompanying motor responses (Fig. 9). Comparable results have been reported previously (Sakimoto and Sakata, 2015).

### Main changes in LFPs recorded in this study in relation to collected GO and noGO correct and incorrect responses

Results collected from the spectral analysis of LFPs recorded in the eight different areas included in this study (Figs. 5-6) indicate that the acquisition of this GO/noGO task required the participation of a distributed network of brain structures and the presence of different oscillatory patters for the proper communication between selected brain circuits. In addition, significant differences were found not only between GO vs. noGO correct and incorrect responses, but also between GO and noGO correct vs. incorrect responses, posing some interesting questions regarding how the experimental animal understands this rather complex task. We should consider that LFP epochs analyzed in this study were restricted to the moment when the animal decides which is the correct response to the stimulus presented on the iPad screen.

On the whole, the significant changes in the dominant peak frequency in relation to the four different experimental responses presented by the PrL cortex, CA1, MC1, and DLS (Fig. 7) could be tentatively related to the proper motor performance of the selected behavior in accordance with the presented visual stimulus and the animal’s decision, a design not directly related to the other brain sites (NAc, BLA, MD, and DMS) included in this study. In this situation, the PrL area would play a double (cognitive vs. behavioral selection) role in this type of associative learning task.

The set of spectrograms illustrated in Figs. 8 and 9 clearly separate cognitive from behavioral information processing, particularly for the PrL cortex, while other brain areas (CA1, MC1, DLS) were more directly involved in programming and performing the necessary motor sequences to reach the iPad and touch the displayed visual stimulus. The other four areas (NAc, BLA, MD, and DMS) presented similar levels of spectral powers across the 2-second epochs, with no definite involvement in any of the four experimental situations. Nevertheless, the coherence analysis illustrated in Fig. 10 suggests that response inhibition for noGO correct responses requires a more precise coordination between several brain centers (PrL, NAc, CA1, MC1, and DLS) involved in this type of associative learning task.

Finally, it is important to point out here that the four areas included in the spectral analysis of LFPs recorded during the initial and final parts of sessions carried out before and after the proper acquisition of the GO/noGO task (Fig. 11) presented more significant changes than during the time segments more directly involved in the learning process (Figs. 5 and 6). This is a further indication that the GO/noGO instrumental learning task involved not only a distributed network of brain structures and functional programs, but also an extension to moments not directly corresponding to stimulus presentations and selected responses.

### Conclusions

In summary, a detailed analysis of LFP activities recorded in the PrL cortex and in other cortical (MC1, CA1) and subcortical (NAc, BLA, DMS, DLS, and DM) areas during the acquisition and performance of an operant conditioning GO/noGO task further supports a selective modulation of neural activities during cognitive vs. locomotor or stationary behavioral activities, depending on the learning stage of the experimental animal.

## Abbreviations

AP: anteroposterior from Bregma
BLA: basolateral amygdala
De: depth from brain surface
D: dorsal
DLS: dorsolateral striatum
DMS: dorsomedial striatum
CA1: hippocampal CA1 area
L: left
LFP: local field potential
mPFC: medial prefrontal cortex
MD: mediodorsal thalamic nucleus
NAc: nucleus accumbens septi
PrL: prelimbic
PSD: power spectral density
MC1: primary motor cortex
R: right
V: ventral

## Acknowledgements

Authors wish to thank Pierre Gussani and José A. Santos Naharro for their technical help in the design and construction of the recording setup, and José M. González Martín for his help in surgical procedures. We also thank Mr Roger Churchill for his contribution to the final version of the manuscript.

## Authors’ contributions

AG and JMD-G designed the experiments; CM-R performed the experiments; AG, CM-R, and GGP analyzed the collected data; CA-S and MAM-P designed the experimental protocol; AG and JMD-G wrote the paper. All authors revised the definitive version of the manuscript.

## Data availability

Collected data in support of the findings of this study are available from the corresponding author upon reasonable request.

